# Identification of conserved proteomic networks in neurodegenerative dementia

**DOI:** 10.1101/825802

**Authors:** Vivek Swarup, Timothy S. Chang, Duc M. Duong, Eric B. Dammer, James J. Lah, Erik E.C.B. Johnson, Nicholas T. Seyfried, Allan I. Levey, Daniel H. Geschwind

## Abstract

Data-driven analyses of human brain across neurodegenerative diseases possess the potential for identifying disease-specific and shared biological processes. We integrated functional genomics data from postmortem brain, including label-free quantitative proteomics and RNA-seq based transcriptomics in an unprecedented dataset of over 1000 individuals across 5 cohorts representing Alzheimer’s disease (AD), asymptomatic AD, Progressive Supranuclear Palsy (PSP), and control patients, as a core analysis of the Accelerating Medicines Project – Alzheimer’s Disease (AMP-AD) consortium. We identified conserved, high confidence proteomic changes during the progression of dementias that were absent in other neurodegenerative disorders. We defined early changes in asymptomatic AD cases that included microglial, astrocyte, and immune response modules and later changes related to synaptic processes and mitochondria, many, but not all of which were conserved at the transcriptomic level. This included a novel module C3, which is enriched in MAPK signaling, and only identified in proteomic networks. To understand the relationship of core molecular processes with causal genetic drivers, we identified glial, immune, and cell-cell interaction processes in modules C8 and C10, which were robustly preserved in multiple independent data sets, up-regulated early in the disease course, and enriched in AD common genetic risk. In contrast to AD, PSP genetic risk was enriched in module C1, which represented synaptic processes, clearly demonstrating that despite shared pathology such as synaptic loss and glial inflammatory changes, AD and PSP have distinct causal drivers. These conserved, high confidence proteomic changes enriched in genetic risk represent new targets for drug discovery.

**Highlights:**

- We distinguish robust early and late proteomic changes in AD in multiple cohorts.
- We identify changes in dementias that are not preserved in other neurodegenerative diseases.
- AD genetic risk is enriched in early up-regulated glial-immune modules and PSP in synaptic modules.
- Almost half of the variance in protein expression reflects gene expression, but an equal fraction is post-transcriptional or -translational.

## Introduction

Dementia, defined as a significant deficit in cognitive domains such as memory, language or executive function leading to a loss of independent functions, affects 9% of the population (Langa et al., 2017; McKhann et al., 2011) and exacts a large societal and financial burden (Alzheimer’s Association, 2019). Alzheimer’s disease (AD) is the most common neurodegenerative dementia and is pathologically characterized by the presence of neurofibrillary tangles, consisting of filamentous hyperphosphorylated microtubule associated protein tau (*MAPT*; tau) inclusions, and senile plaques, consisting of Aβ deposits (for a review see – (De Strooper and Karran, 2016; Goate et al., 1991; Lage et al., 2007; Schneider et al., 2009; Vinters, 2015)). Other neurodegenerative conditions related to AD via alterations in proteostasis and the presence of specific pathological aggregates in brain include Progressive Supranuclear Palsy (PSP), Frontotemporal Dementia (FTD), and Parkinson’s disease (PD) (Arendt et al., 2016; Cummings, 2003; Erkkinen et al., 2018; Kovacs, 2015). Although common and rare genetic variants in *MAPT* cause or increase the risk for AD (Desikan et al., 2015), PSP (Chen et al., 2018), FTD (Olszewska et al., 2016), and PD (Chang et al., 2017), risk for these disorders is heterogeneous and complex (Chang et al., 2017; Chen et al., 2018; Ferrari et al., 2014; Im et al., 2015; Karch and Goate, 2015; Klein and Westenberger, 2012; Lambert et al., 2013; Seelaar et al., 2011). So, despite the substantial advances in identifying genetic contributions to dementia disease risk (Im et al., 2015; Karch and Goate, 2015; Klein and Westenberger, 2012; Kunkle et al., 2019; Seelaar et al., 2011) understanding how risk factors converge on molecular pathways in the brain remains a critical gap in our knowledge (Bonham et al., 2018; Parikshak et al., 2015; Yokoyama et al., 2017).

One powerful approach to provide a data-driven framework for connecting genetic risk to molecular processes disrupted in disease is gene network analysis, which leverages gene co-expression to improve mechanistic models of pathophysiology (Parikshak et al., 2015). To accelerate this process, the National Institutes of Health (NIH) developed the Target Identification and Preclinical Validation Project of the Accelerating Medicines Project – Alzheimer’s Disease (AMP-AD) consortium (Hodes and Buckholtz, 2016) whose goal is to integrate high-throughput genomic and molecular data from disease brain within a network driven structure (Hodes and Buckholtz, 2016; Logsdon et al., 2019). Within this context, several recent large-scale RNA sequencing projects have been conducted (Allen et al., 2018; De Jager et al., 2018; Gaiteri et al., 2016; Logsdon et al., 2019; Mostafavi et al., 2018; Readhead et al., 2018; Zhang et al., 2013), identifying transcriptomic networks and splicing events altered in the cerebral cortex from patients with AD and comparing them to the aging brain. These analyses also showed that genetic risk for AD, rather than being widely distributed, was enriched in specific transcriptomic networks. In a complementary approach, studies using proteomics in postmortem brain from AD patients quantified several thousand proteins and identified AD-related transcriptomic networks. But, these studies were each limited to few relatively small cohorts (Johnson et al., 2018; McKenzie et al., 2017; Ping et al., 2018, 2018; Seyfried et al., 2017; Yu et al., 2018; Zhang et al., 2018). Which aspects of these molecular changes are truly generalizable across cohorts and how they relate to other disorders remains to be determined.

Here we have taken an integrative systems biology approach to identify neurodegenerative dementia-associated protein networks, beginning with high-throughput label-free mass spectrometry-based proteomics in the largest number of subjects to date from multiple cohorts of postmortem human AD brain, as a core component of the AMP-AD consortium (Hodes and Buckholtz, 2016). To detect robust disease-associated changes across multiple cohorts, we used the same processing scheme to harmonize the data across studies and applied a consensus network analysis approach to identify conserved, biologically relevant co-expression modules (Langfelder and Horvath, 2008; Langfelder et al., 2013; Swarup et al., 2019). Additionally, we performed analyses to unravel temporal trajectories of distinct biological processes during the course of disease progression by distinguishing early and late disease changes, where the latter may represent changes associated with chronic disease progression, such as reactive changes. We identified several co-expression modules that were correlated with neuropathology and cognitive decline in AD. We validated these findings in an orthogonal published dataset (Ping et al., 2018) and extended our initial findings across multiple neurodegenerative diseases including FTD with TDP-43 (FTD-TDP), PSP with corticobasal degeneration (PSP-CBD), PD, PD with dementia (PD-D), multiple systems atrophy (MSA), and amyotrophic lateral sclerosis (ALS). We found protein co-expression modules that were dysregulated in neurodegenerative dementias (AD, FTD-TDP, PSP-CBD, and PD-D), but not in related neurodegenerative diseases (PD, MSA and ALS) where dementia is not the defining feature. We evaluated concordance of transcriptomic and proteomic signatures in a subset of samples, which represented the largest comparative dataset to date, finding high correspondence overall, as well as proteome specific changes. To understand the relationship of these core molecular processes with causal genetic drivers, we integrated common variants associated with AD risk with co-expression modules and identified specific biological processes enriched in disease-specific risk variants.

## Results

### Datasets used in the study

We performed label-free high-throughput quantitative proteomics by liquid chromatography coupled with tandem mass spectrometry (LC-MS/MS) on five different study cohorts – the Baltimore Longitudinal Study of Aging (BLSA), Adult Changes of Thought (ACT), Mount Sinai Brain Bank (MSBB), Banner Sun Health Research Institute (Banner) and Mayo Clinic Brain Bank (Mayo, Figure 1), totaling 798 samples representing AD, asymptomatic AD (AsymAD), PSP, and controls from prefrontal, parietal, and temporal cortex. While the BLSA cohort comprised of both prefrontal and parietal cortex samples (n=97 samples) (Johnson et al., 2018; Seyfried et al., 2017) and the Mayo cohort comprised of temporal cortex samples (n=196 samples), the remaining samples (ACT - n=65, MSBB - n=251 and Banner - n=189) were from prefrontal cortex (Figure 1). Detailed clinical and neuropathological scores were available for each cohort and were used in downstream analyses (Table S1). In addition, we analyzed RNA-seq on Mayo samples in which proteomics data was generated, to permit direct comparisons of the transcriptomic and proteomic findings in the same dataset (Allen et al., 2016, 2018). As a validation set, we generated quantitative proteomics data from University of Pennsylvania (UPenn) Brain Bank samples – 384 frontal cortical samples from patients with clinically-diagnosed AD, FTD-TDP, PSP-CBD, PD, PD-D, MSA, and ALS (Figure 1).

**Figure 1.**
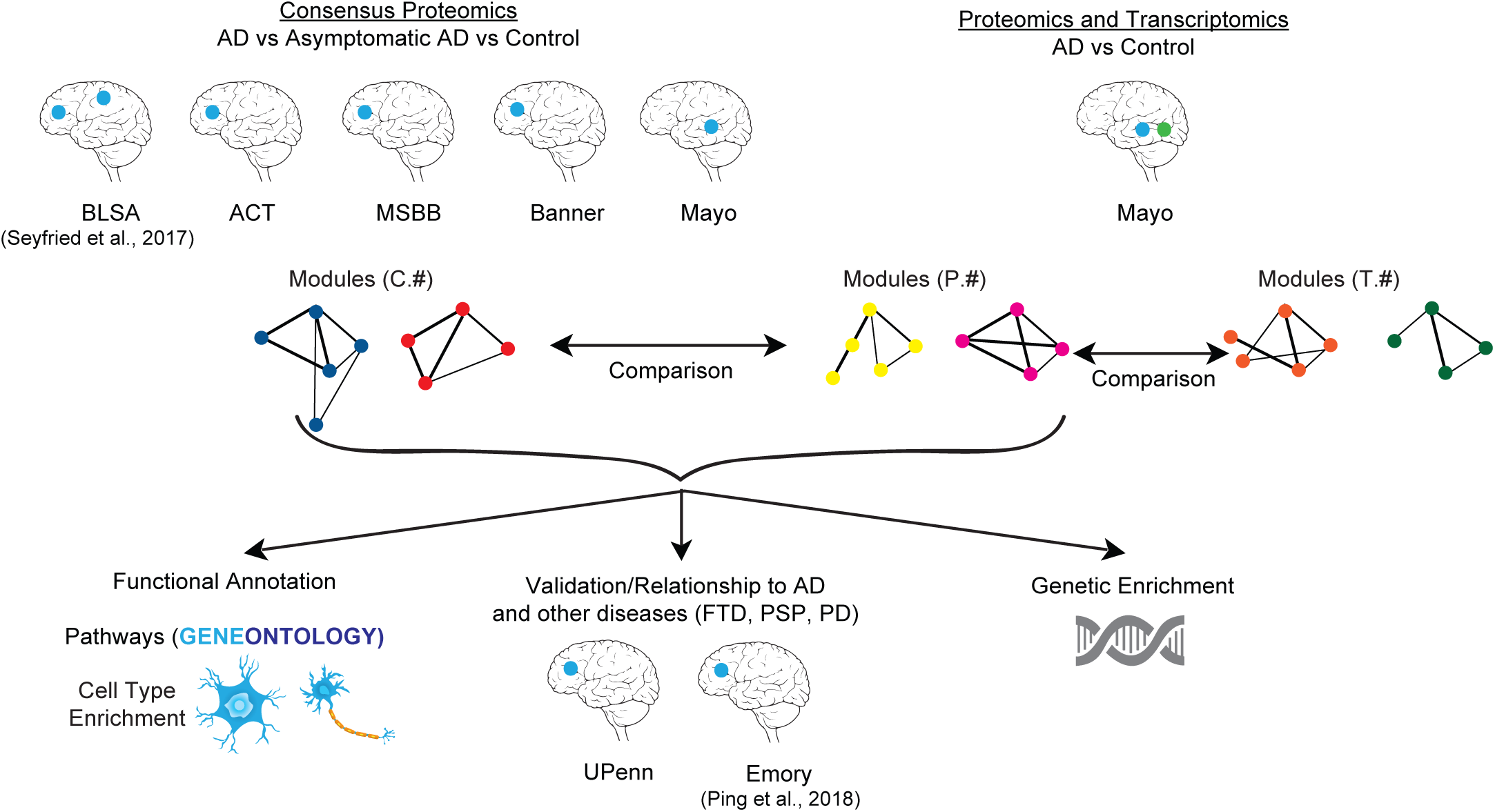
Schematic representation of the various analyses performed in the study Consensus proteomics modules were generated from five different studies – the Baltimore Longitudinal Study of Aging (BLSA), Adult Changes of Thought (ACT), Mount Sinai Brain Bank (MSBB), Banner Sun Health Research Institute (Banner) and Mayo Clinic Brain Bank (Mayo). Furthermore, Mayo dataset had matched samples where proteomics and transcriptomics data were available and thus were analyzed separately. Comparisons were then made between the consensus proteomics modules (labeled with ‘C’), Mayo-only proteomics modules (labeled with ‘P’) and Mayo-matched sample transcriptomics modules (labeled with ‘T’). Functional annotation of the modules was performed using cell-type and gene-ontology. Determining relationship of these modules with other neurodegenerative diseases like frontotemporal dementia (FTD), progressive supranuclear palsy (PSP) and Parkinson’s disease (PD) was performed using completely separate datasets (UPenn, Emory). Common genetic enrichment analysis was used to understand the relationship of modules with causal genetic drivers. The blue dot indicates location of proteomic brain samples. The green dot indicates location of transcriptomic brain samples.

Using a label-free proteomics approach, we were able to identify approximately 65,000 peptides corresponding to approximately 5,000 proteins per sample. To circumvent the problem of missing protein abundance values across different datasets, we set an arbitrary cut-off for a maximum of 40% missing values. In total, 2,005 quantified proteins overlapped in all five datasets (BLSA, ACT, MSBB, Banner, Mayo) and were used for our conservative downstream analysis.

The pathologic hallmarks of Alzheimer’s disease are seen decades before clinical onset of symptoms and are considered the asymptomatic phase of disease (Driscoll et al., 2006; Sperling et al., 2011). The asymptomatic AD and AD samples in our cohort allowed us to profile the molecular alterations that begin early versus late in disease course.

### Identification of robust disease-relevant protein co-expression signature

To place protein expression changes in a systems level framework and find robust co-expression changes irrespective of brain banks and cortical regions, we employed a consensus weighted gene co-expression network analysis (cWGCNA) across the cohorts from the BLSA, ACT, MSBB, Banner and Mayo brain banks (Figure 2A, Figure S1, Table S2). We identified 10 consensus modules (labelled C1-C10) encompassing all four major brain cell types – neurons (C1 and C2 modules), microglia/endothelial cells (C10 module), astrocytes (C8 module), and oligodendrocytes (C5 module) (Figure 2B). We identified three modules (C1, C8 and C10) that were significantly associated with the AD diagnosis (p<0.05) in all five datasets and three additional modules (C2, C3, and C7) significantly correlated with diagnosis (p<0.05) in at least three of the five datasets (see Methods) for a total of six AD associated modules. Three consensus modules (C3, C8, and C10) were upregulated in AD compared to controls, while three consensus modules (C1, C2 and C7) were downregulated in AD (Figure 2B). We leveraged a large cohort of asymptomatic AD cases, where cognition is not affected, but where subjects have sufficient neurofibrillary tangles and plaques for a pathological diagnosis (Hyman et al., 2012), providing a glimpse into early stages of the disease progression. We defined early AD molecular alterations as changes seen in both AsymAD and AD, and late AD molecular alteration as changes only seen in AD. Interestingly, the two glial modules C8 and C10 were upregulated early in the disease, while the remaining modules (C1, C2, C3, and C7) were altered late in disease course (Figure 2B). Table S3 summarizes characteristics of early and late modules associated with AD and their association with other neurodegenerative diseases, analysis of which is further described below. The following website (https://coppolalab.ucla.edu/gclabapps/nb/browser?id=SwarupConsensusProteomicsV4) includes an interactive graphical interface for the consensus proteomic network, allowing exploration of individual proteins and their relationships.

**Figure 2.**
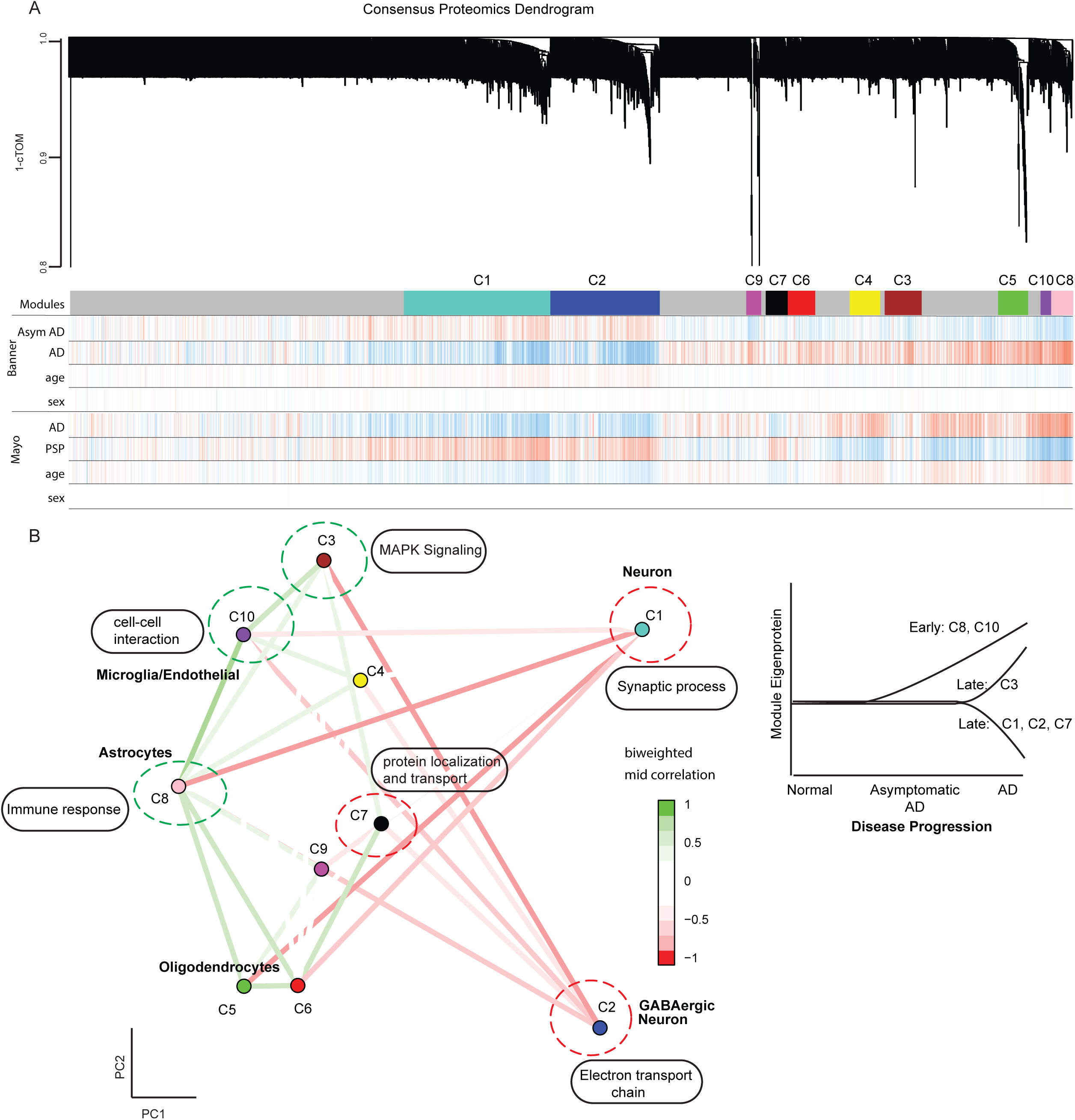
Consensus Proteomics Analyses. (A) Consensus proteomics dendrogram showing the proteomics modules from 2 of 5 different proteomics datasets (Banner and Mayo). Color bars below the modules give information on correlation of disease condition (AD, asymptomatic AD, PSP) and biological covariates (age and gender) with the expression of a particular gene. Red is positive correlation, and blue is anti-correlation. (B) Multidimensional scaling plot on the left demonstrates relationship between consensus proteomics modules and clustering by cell-type relationship. Also shown are the major GO enrichment terms for that module. Modules upregulated in AD (green dotted circle) and downregulated in AD (red dotted circle) are shown. Diagrammatic representation of the trajectory of module eigenprotein with the progression of the Alzheimer’s disease is shown highlighting the early and late proteomic modules identified in the study. cTOM=consensus Topological Overlap Matrix, AD=Alzheimer’s disease, AsymAD = Asymptomatic AD, PSP = Progressive Supranuclear Palsy

### Identification of early changes in the AD proteome

In an attempt to find drivers of disease pathology, we focused our attention on modules significantly correlated with asymptomatic AD cases, representing the earliest disease phase in these samples. This highlighted the C8 and C10 modules, which were significantly upregulated in both AsymAD and AD cases (Figure 3A-J). Gene ontology (GO) analyses indicated that the C8 module, which largely represented genes highly expressed in astrocytes (enrichment p-value=8.9e-7), was enriched for immune response and cell-cell adhesion, consistent with its enrichment in astrocyte marker proteins, such as GFAP and S100B. One hub of the C8 module, MSN, is an essential protein involved in P2X7 receptor-mediated cleavage of amyloid precursor protein (APP) (Darmellah et al., 2012). Another hub protein, PRDX6, is a member of a ubiquitous family of antioxidant enzymes that controls peroxide levels and was reported to be upregulated in astrocytes from patients with Lewy Body Dementia (Power et al., 2002) (Figure 3A). In contrast, the C10 module was enriched in microglial and endothelial markers and showed GO enrichment for cell-cell interactions. One of the hubs of the C10 module, ANXA1, is a central player in the anti-inflammatory and neuroprotective role of microglia, and the clearance of Aβ (McArthur et al., 2010; Park et al., 2017; Ries et al., 2016). Additionally, CLIC1, a C10 module protein hub, is a chloride intracellular channel expressed in microglia and involved in mediating Aβ neurotoxicity (Milton et al., 2008; Skaper et al., 2013) (Figure 3B). This analysis clearly demonstrates that both astroglial and microglial up-regulation are early components of AD, and identifies core genes involved in this process, many of which have been associated with dementia pathophysiology.

**Figure 3.**
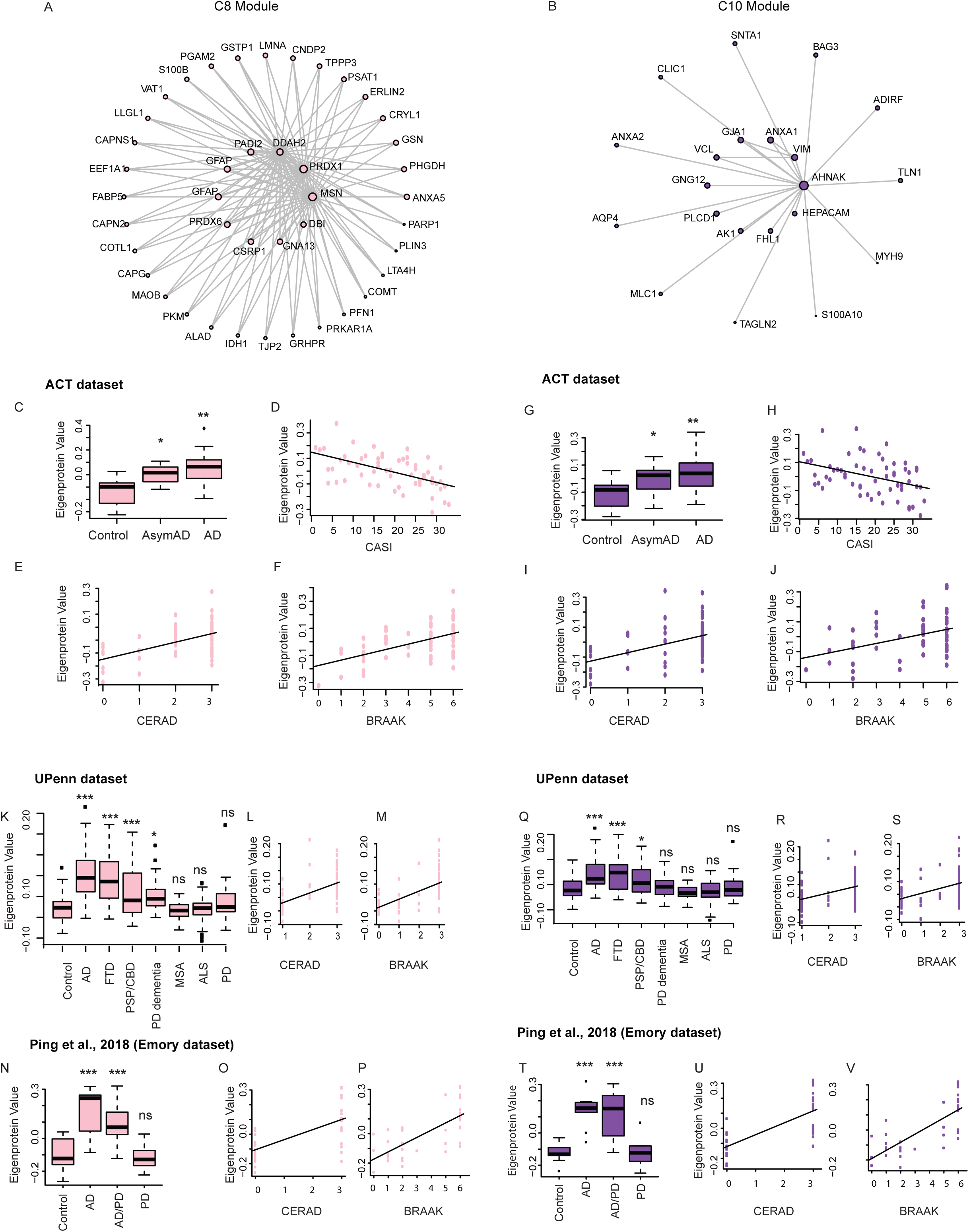
Early proteomic changes in AD. (A, B) Co-expression plot of the consensus proteomic C8 module (A) and C10 module (B) showing hub genes in the center of the module. (C-F) Plots showing C8 module eigenprotein trajectory with diagnosis (C), CASI (D), CERAD (E) and BRAAK scores (F) in ACT dataset. (G-J) Plots showing C10 module eigenprotein trajectory with diagnosis (G), CASI (H), CERAD (I) and BRAAK scores (J) in ACT dataset. (K-M) Using the module protein definitions of the C8 module, the C8 eigenprotein was compared across different diseases in the UPenn dataset (K). CERAD (L) and BRAAK (M) scores for AD samples in the UPenn dataset are shown. (N-P) Validation of C8 module trajectory using Emory dataset (Ping et al., 2018) showing the eigenprotein trajectory with diagnosis (N), CERAD (O) and BRAAK scores (P). (Q-S) Using the module protein definitions of the C10 module, the C10 eigenprotein was compared across different diseases in the UPenn dataset (Q). CERAD (R) and BRAAK (S) scores for AD samples in UPenn dataset are shown. (T-V) Validation of C10 module trajectory using Emory dataset (Ping et al., 2018) showing the eigenprotein trajectory with diagnosis (T), CERAD (U) and BRAAK scores (V). AD=Alzheimer’s disease, AsymAD=asymptomatic AD, FTD=Frontotemporal Dementia, PSP-CBD=Progressive Supranuclear Palsy with Corticobasal Degeneration, PD=Parkinson’s disease, MSA=multiple systems atrophy, ALS=amyotrophic lateral sclerosis. *p<0.05; **p<0.01; ***p<0.005; n.s.=non-significant.

To identify which, if any, of these molecular traits were related to disease progression, we next assessed their relationship to clinical-pathological variables. We found that the C8 module eigenprotein was positively correlated to the hallmark neuropathological traits – amyloid plaques (Consortium to Establish a Registry for Alzheimer’s Disease (CERAD) score (Mirra et al., 1991) – ACT data, ρ=0.52, p=2.9e-5) and neurofibrillary tangles (Braak score (Braak and Braak, 1991) – ACT data, ρ=0.55, p=7.7e-6) – and anti-correlated with cognitive status (Cognitive Abilities Screening Instrument (CASI) score (Teng et al., 1994) – ACT data, ρ= - 0.54, p=1.3e-6) across control, AsymAD, and AD cases (Figure 3E, F). A similar pattern was observed for the C10 module, whose module eigenprotein was positively correlated with CERAD (ACT data, ρ=0.45, p=2.9e-5) and Braak scores (ρ=0.52, p=2.9e-5, Figure 3I, J), and anti-correlated with cognitive status (CASI score – ACT data, ρ= −0.48, p=1.3e-6). Similar trends were observed with the four other study cohorts (Figure S2).

We next analyzed a distinct independent dataset (UPenn; https://doi.org/10.7303/syn5477237; Methods), which in addition to permitting direct independent validation of our findings in AD, also contained seven other neurodegenerative diseases for comparison. Both the C8 and C10 modules were significantly upregulated in AD samples and showed similar trends of association with CERAD and Braak scores in the UPenn dataset (Figure 3K-M, 3Q-S). Interestingly, these early disease-associated modules were upregulated only in neurodegenerative dementias (C8: FTD-TDP, PSP-CBD, PD-D) (Figure 3K) (C10: FTD-TDP, PSP-CBD) (Figure 3Q), but not in other neurodegenerative disease where dementia is not a primary feature (MSA, ALS, PD). We validated our findings using data from a second independent study (Emory (Ping et al., 2018)) that used multiplex isobaric tandem mass tags (TMT) and off-line fractionation for protein quantification method that identified more than 11,000 quantified proteins. In Ping et al. (Ping et al., 2018), the C8 and C10 modules were upregulated in AD and in cases with combined Alzheimer’s disease-Parkinson’s disease pathology, further validating the initial findings (Figure 3N-P, 3T-V).

### Identification of late changes in the AD proteome

We next evaluated proteomic changes occurring in the later, symptomatic phases of AD. We identified one upregulated (C3) and three downregulated (C1, C2 and C7) modules that were significantly correlated with AD diagnosis, but did not change in AsymAD cases. The upregulated module, C3, was enriched with genes involved in MAPK signaling; hub genes of the C3 module included MAPK3 and MAPK1. The C3 module eigenprotein was positively correlated with CERAD and Braak scores and anti-correlated with CASI scores (Figure 4Q-U).

**Figure 4.**
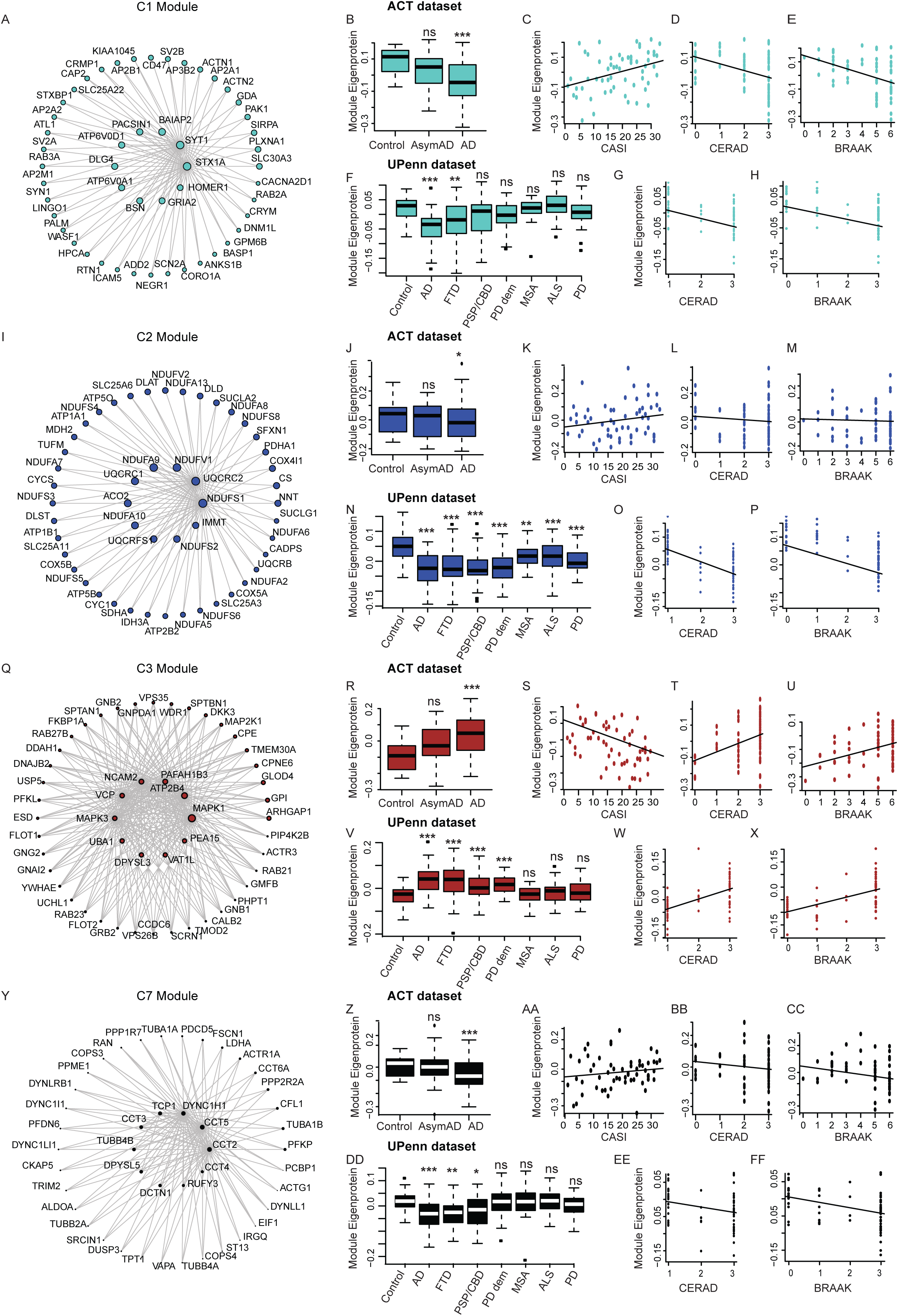
Late proteomic changes in AD. (A) Co-expression plot of the C1 module showing the hub proteins in the center. (B-E) Plots showing C1 module eigenprotein trajectory with diagnosis (B), CASI (C), CERAD (D) and BRAAK scores (E) in ACT dataset. (F-H) Plots showing C1 module eigenprotein trajectory with diagnosis (F), CERAD (G) and BRAAK (H) scores in validation UPenn dataset. (I) Co-expression plot of the C2 module showing the hub proteins in the center. (J-M) Plots showing C2 module eigenprotein trajectory with diagnosis (J), CASI (K), CERAD (L) and BRAAK (M) scores in ACT dataset. (N-P) Plots showing C2 module eigenprotein trajectory with diagnosis (N), CERAD (O) and BRAAK (P) scores in validation UPenn dataset. (Q) Co-expression plot of the C3 module showing the hub proteins in the center. (R-U) Plots showing C3 module eigenprotein trajectory with diagnosis (R), CASI (S), CERAD (T) and BRAAK (U) scores in ACT dataset. (V-X) Plots showing C3 module eigenprotein trajectory with diagnosis (V), CERAD (W) and BRAAK (X) scores in validation UPenn dataset. (Y) Co-expression plot of the C7 module showing the hub proteins in the center. (Z-CC) Plots showing C7 module eigenprotein trajectory with diagnosis (Z), CASI (AA), CERAD (BB) and BRAAK (CC) scores in ACT dataset. (DD-FF) Plots showing C7 module eigenprotein trajectory with diagnosis (DD), CERAD (EE) and BRAAK (FF) scores in validation UPenn dataset. AD=Alzheimer’s disease, AsymAD= asymptomatic AD, FTD=Frontotemporal Dementia, PSP-CBD=Progressive Supranuclear Palsy with Corticobasal Degeneration, PD=Parkinson’s disease, MSA=multiple systems atrophy, ALS=amyotrophic lateral sclerosis. *p<0.05; **p<0.01; ***p<0.005; n.s.=non-significant

The down-regulated modules, C1 and C2, were enriched in neuronal markers and enriched in GO terms like synaptic processes and neurotransmitter secretion pathways. The downregulation of C1 and C2 modules is consistent with neuronal loss occurring in symptomatic disease stages. Module C1 included hubs consisting of pre-and post-synaptic vesicle proteins such as HOMER1, DLG4, and SYT1 (Figure 4A). The C2 module consisted of mitochondrial electron transport chain subunits (NDUFs, ACO2) and ATPase subunits (ATP1A1, ATP1A2, etc.) (Figure 4I). The downregulated C7 module was enriched in protein localization and transport function as GO terms comprises of several cytoskeletal proteins — dynactins and tubulin subunits (DCTN1, TUBB4B) and molecular chaperones (TCP1, CCT2, CCT3) (Figure 4Y). The module eigenprotein of all three modules (C1, C2 and C7) decreased with progression of disease as measured by CERAD and Braak scores (Figure 4B-E, 4J-M, 4Z-CC).

We next determined whether these late modules could be validated in other data sets, similar to the early modules changing in pre-symptomatic patients. Modules C1, C3, and C7 were preserved and solely associated with AD and other neurodegenerative dementias (C1: FTD-TDP) (Figure 4F-H) (C3: FTD-TDP, PSP-CBD, PD-D) (Figure 4V-X) (C7: FTD-TDP, PSP-CBD) (Figure 4DD-FF), whereas C2 was associated with AD and a broader range of neurodegenerative conditions (FTD-TDP, PSP-CBD, PD-D, MSA, ALS, PD) (Figures 4N-P). Testing for preservation of these modules in the additional dataset from Ping et al. (Ping et al., 2018), revealed that all modules were preserved and associated with AD and cases with combined Alzheimer’s disease and Parkinson’s disease (Supplemental Figures 3-4).

### Comparison of the AD proteome and transcriptome

Most understanding of biological mechanisms occurs at the level of proteins, and yet most “-omics” data have been represented by transcriptomics. We reasoned that it would be important to understand how transcriptomic data was reflected at the level of proteomics collected from the same individuals in the same brain region, especially since little such comparative data is available in dementia (McKenzie et al., 2017; Seyfried et al., 2017). We leveraged 168 neurodegenerative disease and 28 control cortical samples in the Mayo cohort from which proteomics and RNA-seq data was generated from the same samples, making it the largest such comparison to date.

We created separate co-expression networks for RNA (labelled T1-T21 modules from 11,140 transcripts) and protein (labelled P1-P15 modules from 3,849 proteins, Figure S5). As expected, the protein modules overlapped significantly with the consensus proteomic modules identified in the consensus analysis (Figure 5A; Figure S6). We found that the glial consensus modules upregulated early in AD, C8 (astrocyte, immune process) and C10 (microglia/endothelial cells, cell-cell adhesion), overlapped with the Mayo protein P3 module (C8-P3 hypergeometric overlap OR=25; C10-P3 hypergeometric overlap OR=33). This P3 module enriched for astrocyte and microglial cell types and also overlapped with multiple Mayo transcriptomic modules that were enriched for astrocyte and microglial cell types – T3, T8, T9, T18 and T21 (Figure 5A; hypergeometric overlap OR ranges 6-12).

**Figure 5.**
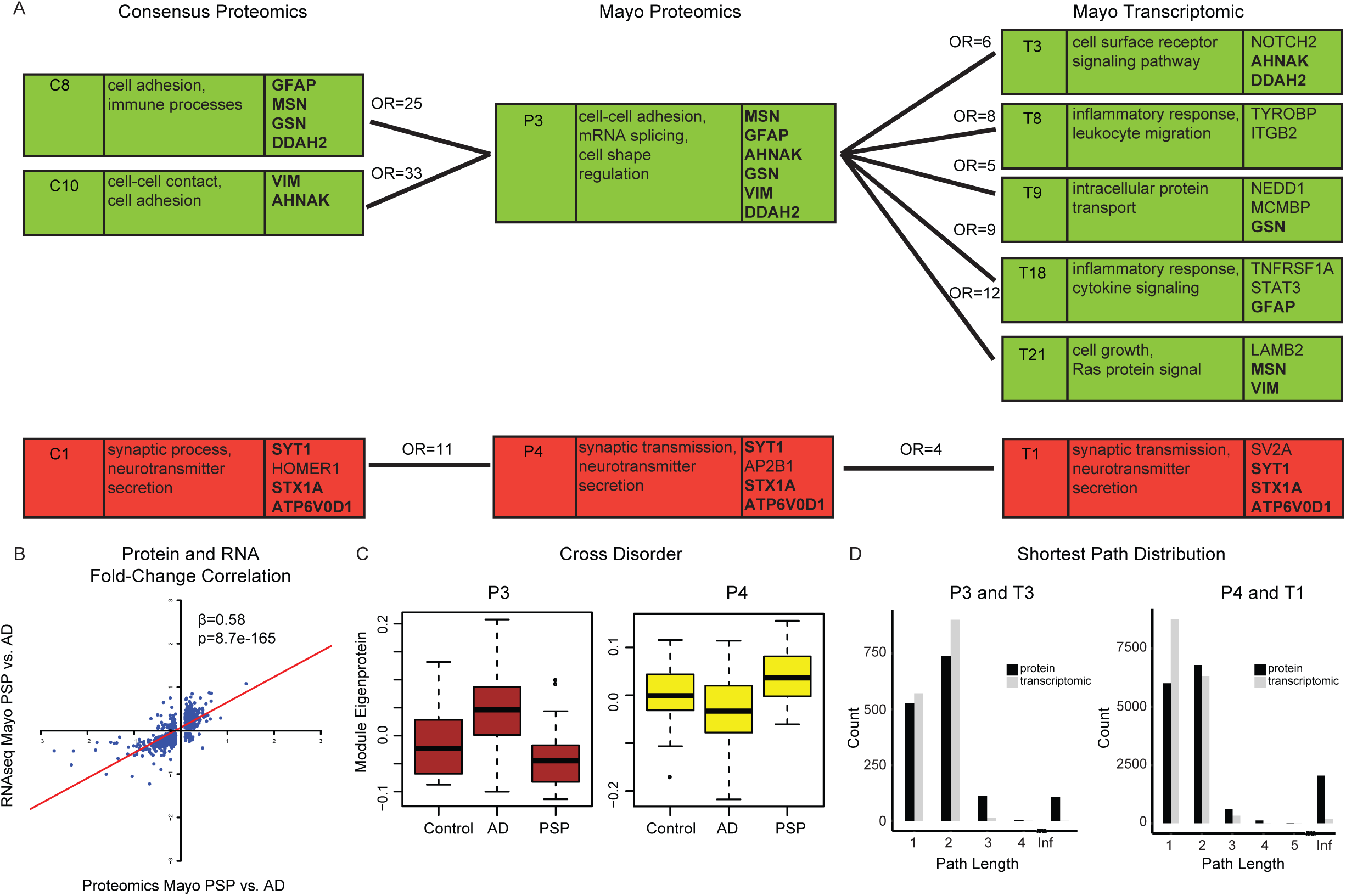
Proteomics and Transcriptomic Analyses. (A) Relationship of Consensus Proteomic, Mayo Proteomics and Mayo Transcriptomic Modules. The first column is the module name; the second column includes Gene Ontology terms; and the third column includes genes that are hubs within the module. Modules with significant hypergeometric overlap between Consensus Proteomics and Mayo Proteomics, and between Mayo Proteomics and Mayo Transcriptomics are shown. Numbers above connecting lines are the odds ratio of hypergeometric overlap. These modules were also upregulated (green) or downregulated (red) in Alzheimer’s disease compared to controls or Progressive Supranuclear Palsy. (B) For differentially expressed genes in PSP compared to AD (FDR<0.1), the PSP to AD log2 fold-change at the RNA level correlated with the fold-change at the protein level. (C) Module eigenprotein trajectory with disease for P3 and P4 Mayo proteomic modules. (D) Distribution of shortest path between pairs of genes for P3 and T3 modules, and P4 and T1 modules. Infinite distance indicates unconnected gene pairs.

Identification of modules that were preserved at the proteomic and transcriptomic levels (Methods, Figure 3 and 5A) allowed us to focus on shared, high-confidence biological processes and specific hub genes associated with disease. These genes include MSN (modules C8, P3, T21, Figure 3A), GFAP (modules C8, P3, T8, Figure 3A), DDAH2 (modules C8, P3, T3, Figure 3A), VIM (modules C10, P3, T21, Figure 3B), and AHNAK (modules C10, P1, T3, Figure 3B). While MSN, a microglial enriched gene, is involved in cleavage of APP (Darmellah et al., 2012), GFAP and VIM are reactive astrocyte markers (Kamphuis et al., 2012), reinforcing the role of glia in early AD pathogenesis (Dzamba et al., 2016; Efthymiou and Goate, 2017; Garwood et al., 2015; Heneka et al., 2015; Nagele et al., 2004; Wang et al., 2015). DDAH2 is involved in nitric oxide production, a component of the immune response (Lambden et al., 2016). A missense variant in AHNAK was recently associated with AD in a large exome array study (Sims et al., 2017)

The consensus module downregulated in AD (C1) overlapped with the Mayo protein module P4 (OR=11) and T1 transcriptomic module (OR=4) (Figure 5A). These modules were enriched in GO terms like synaptic processes and neurotransmitter secretion pathways, and neuronal cell type. Shared hubs among all overlapping modules (C1, P4 and T1) included synaptic proteins like SYT1 and STX1A, both involved in vesicle trafficking. Studies have shown SYT1 interacts with the APP ectodomain to promote Aβ generation (Gautam et al., 2015) and STX1A binds another AD gene presenilin 1 (Smith et al., 2000). Shared hub ATP6V0D1 is a subunit of vacuolar-ATPase involved in lysosomal acidification (Colacurcio and Nixon, 2016) and has been linked to AD pathogenesis.

The availability of transcriptomics and proteomics datasets from the same samples in the Mayo dataset (Control – n=27, PSP – n=80 and AD – n=80 samples) allowed us to compare proteomic and transcriptomic changes at the gene level in two common tauopathies, AD and PSP. For differentially expressed genes in PSP compared to AD (FDR<0.1), the PSP to AD fold-change at the RNA level was significantly correlated with the fold-change at the protein level (β=0.58, R^2^=0.56, p=8.7e-165, Figure 5B). This indicates that slightly more than half of the variance in protein expression reflects gene expression, but that an almost equivalent proportion of the signal is post-transcriptional, strongly supporting the utility of protein profiling and the complementarity of these two data sets.

Genes upregulated in AD compared to PSP at both the RNA and protein levels included GO terms associated with cell-cell adhesion and nuclear-transcribed mRNA catabolic process, similar to GO terms for modules significantly upregulated in AD (C8, C10, P3, T3) (Figure 3A, 3G, 5A, 5C). On the other hand, downregulated genes in AD at both the RNA and protein levels included GO terms like neurotransmitter secretion, synaptic vesicle exocytosis, and mitochondrial respiratory chain complex I assembly, which were similar to modules significantly downregulated in AD (C1, P4 and T1) (Figure 4A, 5A, 5C).

In addition to comparing Mayo proteomic and transcriptomic module gene membership and overlap, we compared module connectivity of overlapping modules in an attempt to parse any structural differences between proteomic and transcriptomic networks. We calculated the shortest path between all pairs of genes in a module and found that the proteomic modules, in general, had lower connectivity than the transcriptomic modules (Figure 5D, Table S4), which was not surprising given the lower total number of proteins quantified compared to transcriptomics. We also calculated the clustering coefficient, which measures the connectivity among a gene’s neighbors and higher values signify greater connectivity. The clustering coefficient was significantly larger in transcriptomic modules compared to proteomic modules (Table S4, p-value range <2.2e-16 to 7.2e-4), consistent with tighter connections between transcripts represented within transcriptomic modules. As the total number of proteins measured for the Mayo proteome was smaller than the number of transcripts measures for the Mayo transcriptome, proteomic modules may be missing hub genes due to lack of measurement, as module path length increases if hub genes are removed (e.g. (Chandran et al., 2016)).

The electronic transport chain (C2), protein transport (C7), and MAPK (C3) late proteomic modules did not overlap with transcriptomic modules in our study, suggesting unique disease signatures could be identified via co-expression proteomic analyses (Seyfried et al., 2017). To investigate why these proteomic modules did not overlap with transcriptomic modules, we found that a large percentage of genes from these non-overlapping proteomic modules were not assigned to any transcriptomic modules (65 of 219 for C2, 6 of 44 for C7 and 26 of 74 for C3). This indicates that the RNA of these unassigned genes, while expressed at moderate levels, did not have strong co-expression with genes in any transcriptomic module. By measuring protein we were able to detect co-expression, which likely reflects post-transcriptional regulation, as it was not captured in the transcriptome. Consistent with this, we observe that the PSP to AD RNA fold-change was correlated with the protein fold-change for all differentially expressed genes as described above (β=0.58), whereas the RNA to protein fold-change was less correlated or negatively correlated for these non-overlapping proteomic modules (β= −0.41 for C2, β=0.04 for C7, and β=0.30 for C3). Overall, the non-overlapping proteomic modules represented genes with significantly lower RNA to protein correlation than the average gene, likely due to a combination of post-transcriptional and post-translational regulation (Filipowicz et al., 2008; Glisovic et al., 2008; Hess and Stamler, 2012; Hicke, 2001; Vosseller et al., 2002; Zhao et al., 2017). For example, the MAPK pathway (C3) protein levels are regulated post-transcriptionally via RNA binding proteins and microRNA (Sugiura et al., 2011; Whelan et al., 2012), and post-translationally, via ubiquination (Lin et al., 2003).

Previous studies have associated both proteomic and transcriptomic mitochondrial modules with AD (Johnson et al., 2018; Kim and Choi, 2015; Miller et al., 2008; Mostafavi et al., 2018; Munoz and Ammit, 2010; Ping et al., 2018; Shih et al., 2015). Protein transport and localization module association with AD are unique to proteomic studies (Agostinho et al., 2015; Culvenor et al., 1997; Schubert et al., 1991), rather than transcriptomics studies. The module enriched in MAPK signaling identified robustly here has not been previously associated with AD in either proteomic or transcriptomic studies. MAPK signaling is involved in inflammatory responsive pathways (Jones and Kounatidis, 2017; Lee and Kim, 2017; Zhu et al., 2002) and is induced by interleukins, Aβ, excitotoxicity, and oxidative stress (Munoz and Ammit, 2010; Shih et al., 2015). The MAPK signaling family is also involved in tau hyperphosphorylation (Guise et al., 2001; Kim and Choi, 2010; Wang and Liu, 2008) and hippocampal memory formation via the ERK pathway (Kelleher et al., 2004). C3 module hub proteins include MAPK1, the gene encoding ERK2, MAPK3, the gene encoding ERK1, and PEA-15, which was found to activate ERK1/2 in a Ras-dependent manner (Ramos et al., 2000). The potential contribution of PEA-15 to AD pathogenesis has not been studied. Thus, proteomic co-expression analysis can identify unique molecular disease processes previously associated with AD, as well as potentially novel mechanisms, and as such represents an important complement to transcriptomic analyses.

### Assessment of Genetic Risk within Proteomics Modules

The observed proteomic changes may be a cause or consequence of the disease. Since genetic variation associated with disease is causal in this regard, we next integrated genome-wide association studies (GWAS), so as to understand each module’s relationship to potential causal mechanisms (Gandal et al., 2018a; Parikshak et al., 2016). We used summary data from AD GWAS as input for MAGMA (de Leeuw et al., 2015), which controls for confounders including gene-length and GC content, to generate a single p-value for each protein coding risk loci. After correcting for multiple comparisons, common variants from AD GWAS (Lambert et al., 2013) were significantly enriched in the consensus C8 (astrocyte) and C10 (microglia) module (Figure 6A), in addition to the overlapping Mayo proteomics P3 module (Figure 6B) and Mayo transcriptomics T3 and T8 modules (Figure 6C). This is consistent with previous reports of AD candidate gene enrichment in glial modules (Seyfried et al., 2017; Zhang et al., 2013). Combined with the early up-regulation of C8 and C10, and their preservation in multiple data sets, this highlights the C8 and C10 co-regulated set of genes as potential targets for disease modification. In contrast, common variants from PSP GWAS (Höglinger et al., 2011) were significantly enriched in the neuronal, C1 module (Figure 6A), in addition to the overlapping Mayo proteomics P4 module (Figure 6B) and Mayo transcriptomic, T1, module (Figure 6C), indicating that causal risk variants in these two tauopathies associated with dementia (AD and PSP) differ in terms of the specific cell types that they causally impact (Figure 6D).

**Figure 6.**
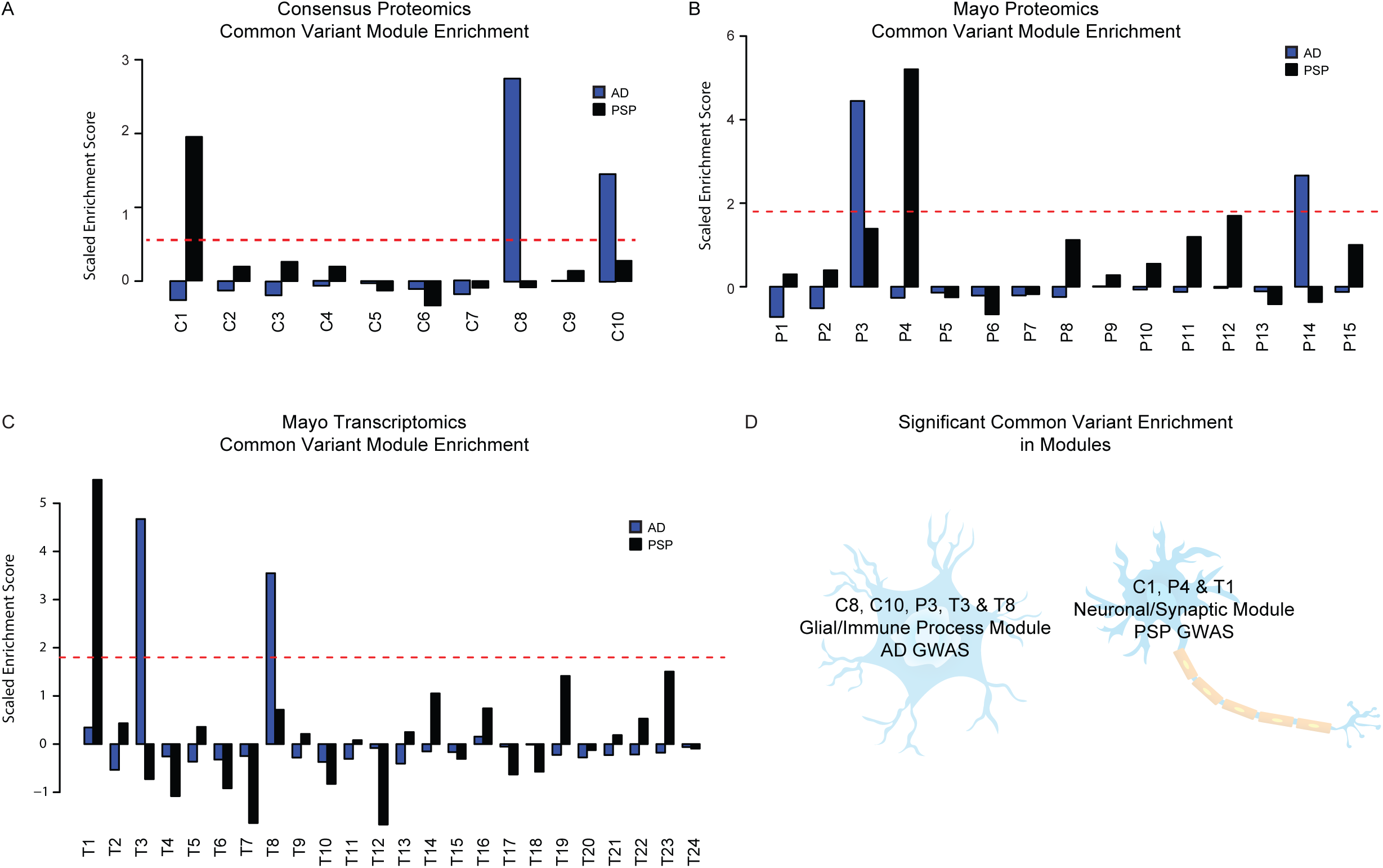
Common variant enrichment in consensus proteomics modules. (A) Mean scaled enrichment of GWAS hits from AD (Lambert et al 2011) and PSP (Höglinger et al., 2011) in consensus proteomics modules, (B) Mayo proteomic modules, and (C) Mayo transcriptomic modules. (D) Diagram showing enrichment of AD and PSP common variants from GWAS studies in significant consensus proteomic modules, Mayo proteomic modules, and Mayo transcriptomic modules, in addition to their cell-type enrichment.

## Discussion

Quantifying changes in the brain proteome provides the most direct way of understanding proteinopathies like Alzheimer’s disease (AD). Understanding the key drivers in a proteomic network has direct implications in identifying druggable protein targets for the disease. With this in mind, we embarked on creating and analyzing the largest proteomic dataset of postmortem brain from patients with various stages of AD to date. By combining multiple cohorts of postmortem human AD brains with high-throughput label-free proteomics, we detected robust and reproducible proteomic modules representing early and late changes associated with AD. We found distinct proteomic modules enriched for AD, as opposed to PSP genetic risk variants, which were dysregulated in orthogonal datasets of AD and other neurodegenerative disease, and conserved at the transcriptomic levels. This analysis provides new insights into the pathogenic processes of AD that are conserved among neurodegenerative dementias, but have disease specificity at the causal genetic level.

One of the major challenges in using proteomics approaches is the limited scalability and reproducibility of proteomic studies to identify disease-specific alterations in large cohorts. With the use of label-free quantitative mass spectrometry, researchers are beginning to unravel proteomic changes in AD (Andreev et al., 2012; Neilson et al., 2011; Seyfried et al., 2017); however scalability and reproducibility between cohorts remains a major concern (Tabb et al., 2010). While previous AD proteomic studies incorporated one or two datasets (Johnson et al., 2018; McKenzie et al., 2017; Ping et al., 2018; Seyfried et al., 2017; Yu et al., 2018; Zhang et al., 2018), we integrated data representing five different cohorts and used a bioinformatics approach to define robust disease-associated proteomic networks. Our consensus WGCNA approach bypassed the need for batch correction, which can lead to the removal or compression of the disease signal (Gandal et al., 2018b; Swarup et al., 2019).

The early molecular changes in sporadic AD disease pathogenesis, particularly in asymptomatic AD, are not well characterized (Driscoll et al., 2006; Hohman et al., 2016). Previous proteomic studies identified inflammatory proteomic modules enriched in glial markers upregulated in asymptomatic AD (Andreev et al., 2012; Johnson et al., 2018; Seyfried et al., 2017). Our larger dataset confirmed that the glial modules – the C8 astrocyte and C10 microglia modules were reproducibly and reliably upregulated early in asymptomatic AD. Interestingly, AD genetic risk variants identified in GWAS were enriched in the C8 astrocytic and C10 microglia modules, providing evidence that genetic causal drivers for early stage AD are associated with glia and thus primarily impact glial biology. While increasing evidence demonstrates the role of glial cells in AD, (Dzamba et al., 2016; Efthymiou and Goate, 2017; Heneka et al., 2015; Nagele et al., 2004; Wang et al., 2015) our study indicates that dysregulation in astrocytes and microglia is an early molecular change in the disease.

Similar to our study, previous proteomic studies (Andreev et al., 2012; Johnson et al., 2018; Seyfried et al., 2017; Zhang et al., 2018) identified neuronal (DeKosky and Scheff, 1990; Selkoe, 2002; Terry et al., 1991), protein transport (Agostinho et al., 2015; Culvenor et al., 1997; Schubert et al., 1991), and mitochondrial (Cadonic et al., 2016; Eckert et al., 2011; Moreira et al., 2010; Swerdlow, 2018) signatures downregulated later in AD progression. Although single cohort studies showed decreased cytoskeletal and microtubule signatures in AD (Andreev et al., 2012; Seyfried et al., 2017; Zhang et al., 2018), our study’s microtubule C5 module was not consistently downregulated in AD across cohorts.

Like AD proteomic analyses, transcriptomic analyses also showed upregulation of inflammatory astrocytic and microglial signatures and downregulation of neuronal signatures (Mostafavi et al., 2018; Seyfried et al., 2017; Swarup et al., 2019; Wang et al., 2016). One transcriptomic study identified a mitochondrial module positively associated with AD (Mostafavi et al., 2018), which contrasted with our and other studies’ (Johnson et al., 2018; Ping et al., 2018) proteomic mitochondrial modules that were negatively associated with AD. However, our consensus proteomic analyses did not identify cell cycle, chromatin modification, glucuronosyltransferase, or axon guidance modules that were identified in transcriptomic studies (Mostafavi et al., 2018; Wang et al., 2016; Zhang et al., 2013), perhaps in part, a limitation of proteomic studies, where less soluble or very large membrane associated proteins are more difficult to detect. This again supports the complementarity of multi-omics approaches and the value of measurements of distinct molecular moeties.

Co-expression proteomic analyses yielded protein specific findings not seen in transcriptomic studies. The consensus proteomic electronic transport chain (C2), protein transport (C7), and MAPK (C3) modules did not overlap with transcriptomic modules. Consistent with this representing differences at the post-transcriptional and post-translational levels that cannot be identified by measuring the transcriptome alone, we find that these genes show lower protein-RNA correlation compared with the average observed across the genome. As late biological processes, these may be adaptive responses to AD pathology or propagate disease progression after AD pathology is established. Mitochondrial modules were associated with AD using transcriptomics and proteomics in previous studies (Johnson et al., 2018; Kim and Choi, 2015; Mostafavi et al., 2018; Munoz and Ammit, 2010; Ping et al., 2018; Shih et al., 2015). However, protein transport modules were only seen in previous proteomic co-expression studies (Agostinho et al., 2015; Culvenor et al., 1997; Schubert et al., 1991) and a MAPK module has not been described in proteomic or transcriptomic studies.

The AD-associated consensus proteomic modules created using five different cohorts were highly preserved not only in a published dataset employing a different TMT-labelled proteomic technique (TMT-LC/MS-MS) (Ping et al., 2018), but also in a large unpublished dataset (UPenn) comprised of several different neurodegenerative diseases, which allowed us to compare these robust changes observed in AD, in other neurodegenerative disorders. This analysis demonstrated modules were shared across disorders, implicating potentially convergent processes at some stage of disease. However, individual disorders had unique module genetic enrichments, suggesting distinct causal mechanisms. The astrocytic C8 module was upregulated in the AD, FTD-TDP, PSP-CBD, and PD-D, and the microglial C10 module was upregulated in AD, FTD-TDP, PSP-CBD. Although this highlights the role of astrocyte (González-Reyes et al., 2017; Perez-Nievas and Serrano-Pozo, 2018; Radford et al., 2015) and microglial (Bachiller et al., 2018; Fernández-Botrán et al., 2011; Hansen et al., 2018; Ishizawa and Dickson, 2001; Navarro et al., 2018; Radford et al., 2015) up-regulation in the pathogenesis of these diseases, their time course and the genetic enrichment analysis shows that their causal role in non-AD dementias is likely different from AD. For example, the C8 and C10 modules’ early up-regulation in AD coupled with the enrichment of common genetic risk from AD, but not PSP, supports its role as a causal driver in AD, but not PSP. Similarly, synaptic loss is a common theme across all neurodegenerative disease, but genetic enrichment in specific synaptic modules is not observed universally. The downregulated C1 module was dysregulated exclusively in neurodegenerative dementia cases (AD, FTD-TDP), and not other neurodegenerative conditions suggesting that this module may contribute to shared pathogenic pathways related to cerebral cortical dysfunction (Bereczki et al., 2018; Bigio et al., 2001; Clare et al., 2010; Hassan et al., 2011). Interestingly, this synaptic C1 module was enriched for common genetic variants in PSP, indicating that these synaptic variants may be causal drivers of PSP, but not AD.

Our study is the largest to comprehensively compare RNA and protein co-expression in dementia. At the gene level, we showed a high correlation of RNA and protein differential expression in PSP compared to AD. At the co-expression network level, glial early proteomic modules (C8, C10) and one synaptic late proteomic module (C1) significantly overlapped with transcriptomic modules, suggesting these biological processes are altered at both the RNA and protein level. As in the previous study investigating RNA and protein module overlap (Seyfried et al., 2017), overlapping modules have shared cell-type markers. The concurrence between proteomic and transcriptomic modules allowed identification of preserved hub genes.

The C8 and C10 modules showed transcriptomic module concordance with preserved hub genes, early up-regulation in AD, convergence with neurodegenerative dementias, and AD causal genetic enrichment. Although we showed proteomic modules have less connectivity compared to transcriptomic modules, proteins represent druggable targets (Accelerating Medicines Partnership in Alzheimer’s Disease (AMP-AD), 2019; Griffith et al., 2013; Hopkins and Groom, 2002; Oprea et al., 2018).

Both GSN and MSN are preserved C8 hub genes linked to AD pathogenic mechanisms. Moesin phosphorylation along with ezrin and radixin, was shown to be required for P2X7R-dependent proteolytic process related to amyloid precursor protein that leads to soluble APP release, which is neuroprotective (Colacurcio and Nixon, 2016; Darmellah et al., 2012). The ezrin/radixin/moesin complex defined an upregulated proteomic module previously associated with AD (Seyfried et al., 2017; Zhang et al., 2018). Moesin is considered druggable based on the Drug Gene Interaction database (Accelerating Medicines Partnership in Alzheimer’s Disease (AMP-AD), 2019; Griffith et al., 2013) and drugs targeting ezrin/radixin/moesin phosphorylation could lead to improved amyloid clearance. In addition, GSN, gelsolin, is known to bind Aβ (Chauhan et al., 1999), inhibit Aβ fibrillization (Ray et al., 2000) and be anti-amyloidogenic in transgenic AD mouse models (Antequera et al., 2009; Hirko et al., 2007; Yang et al., 2014). It is considered druggable (Accelerating Medicines Partnership in Alzheimer’s Disease (AMP-AD), 2019) and histone deacetylase inhibitors can increase gelsolin levels (Hoshikawa et al., 1994; Kamitani et al., 2002), potentially increasing amyloid clearance. Other preserved C8 hub genes such as DDAH2 have not yet been linked to AD pathogenesis. DDAH2 is an enzyme that hydrolyzes asymmetrically methylated arginine residues on proteins, which in turn increases nitric oxide synthase (NOS) (Tran et al., 2000). As an enzyme, DDAH2 is potentially druggable (Accelerating Medicines Partnership in Alzheimer’s Disease (AMP-AD), 2019) and may be associated with AD given the immunoregulatory role of NOS (Lambden et al., 2016), and the function of heterotrimeric S-nitrosylase complex. The heterotrimeric S-nitrosylase complex includes inducible NOS, S100A8 and S100A9, which selectively perform S-nitrosylation on multiple targets including the previously mentioned moesin and ezrin (Jia et al., 2014).

In module C10, two hub genes, ANXA1 and GJA, have been previously associated with AD mechanisms. ANXA1 contributes to microglia-based neuroprotection and Aβ clearance (McArthur et al., 2010; Park et al., 2017; Ries et al., 2016). GJA1 is a connexin hemichannel which has been nominated previously as a target for further study in the AMP-AD based transcriptomic network analyses (Goodenough and Paul, 2003; Kajiwara et al., 2018). Cell biological investigations have shown that Gja1-/- astrocytes co-cultured with neurons improved neuronal survival (Kajiwara et al., 2018) and hemichannel blockers were beneficial in murine AD models (Yi et al., 2017). VIM is a preserved hub gene known to be elevated in reactive astrocyte surrounding plaques (Jing et al., 2007; Pekny and Pekna, 2014). *In vivo* studies have been contradictory as AD mice lacking GFAP and VIM had both increased Aβ in one study (Kraft et al., 2013) and no change in amyloid β staining in another (Kamphuis et al., 2015). These proteins are druggable targets (Accelerating Medicines Partnership in Alzheimer’s Disease (AMP-AD), 2019) and warrant further study.

Our multi-omics, multi-cohort bioinformatics approach provides new insights into early and late changes in AD that are shared with the transcriptomic signature and in other neurodegenerative dementias, but show disease specificity for genetic risk. Shared neurodegenerative dementia modules with consistent molecular alterations at the genetic, transcriptomic, and proteomic levels are ripe for future investigation as drug targets.

## Acknowledgements

Support for this research was provided by funding from the National Institute of Neurological Disorders and Stroke Translational Neuroscience Training Grant (R25NS065723), National Institute on Aging (R01AG053960, R01AG057911, R01AG061800, RF1AG057470, RF1AG05747101), the Accelerating Medicine Partnership for AD (U01AG046161), the Emory Alzheimer’s Disease Research Center (P50AG025688), and the NINDS Emory Neuroscience Core (P30NS055077).

## Author Contributions

Conceptualization, V.S., T.S.C, J.L., E.E.C.B.J., N.T.S., A.I.L., and D.H.G.; Methodology, V.S., T.S.C, D.M.D., E.B.D., N.T.S., A.I.L., and D.H.G.; Software, V.S. and T.S.C; Formal Analysis, V.S., T.S.C., D.M.D., and E.B.D.; Investigation, D.M.D. and E.B.D.; Resources, J.J.L., E.E.C.B.J, N.T.S., A.I.L., and D.H.G; Data Curation, V.S.; Writing – Original Draft, V.S., T.S.C, and D.H.G.; Writing – Review & Editing, V.S., T.S.C., J.L., E.E.C.B.J., N.T.S., A.I.L., and D.H.G.; Visualization, V.S. and T.S.C.; Supervision, N.T.S., A.I.L., and D.H.G.; Funding Acquisition, N.T.S., A.I.L., and D.H.G.

## STAR ★ Methods

### Lead Contact and Material Availability

Further information and requests for resources should be directed to and will be fulfilled by the Lead Contact, Daniel H. Geschwind. This study did not generate new unique reagents.

### Experimental Model and Subject Details

#### Human Samples

Postmortem human brain tissues were obtained from BLSA, ACT, MSSBB, Banner, Mayo, UPenn, and Emory. All the human samples were obtained under their respective institutional review board. For details about sample-level metadata including diagnosis, gender, neuropathological criteria, etc please see Table S1. The BLSA samples consisted of 97 samples from the dorsolateral prefrontal cortex (BA9 area) and precuneus (parietal cortex, BA7) representing 15 controls, 15 AsymAD and 20 AD cases (Seyfried et al., 2017). The ACT samples consisted of 65 samples from 12 controls, 14 AsymAD and 39 AD cases from prefrontal cortex area. The MSBB consisted of 251 prefrontal cortex samples from 78 controls, 28 AD possible, 13 AD probable and 132 AD definite cases based on clinical dementia rating (CDR) scores (Morris, 1993). The Banner samples were also from prefrontal cortex of 30 controls, 28 mild cognitive impairment cases, 33 AsymAD and 98 confirmed AD cases. The Mayo samples were from temporal cortex consisting of 28 controls, 84 AD, and 84 PSP cases. The UPenn samples were from prefrontal cortex consisting of 43 controls, 42 AD, 58 ALS, 29 FTD-TDP, 22 MSA, 33 PD, 58 PD-dementia (PDD) and 48 PSP-CBD. Emory samples were previously published (Ping et al., 2018; Seyfried et al., 2017). This study was performed under the auspices of the UCLA Office of Human Research Protection, which determined that it was exempt because samples were anonymous pathological specimens (IRB# 15-001397).

### Method Details

#### Quantitative Proteomics

Label-free quantitative proteomics were performed at the Emory Proteomics Core, Emory University, USA. Detailed methods were published elsewhere (Seyfried et al., 2017). The label free quantitation (LFQ) algorithm in MaxQuant (Cox and Mann, 2008) was used for protein quantitation. The quantitation method only considered razor and unique peptides for protein level quantitation. The LFQ intensities were log2 transformed for downstream analyses.

#### Covariate correction

Protein LFQ (proteomics) and normalized FPKM values (RNA-seq) were assessed for effects from biological covariates (diagnosis, age, gender) and technical variables (batch, brain bank, etc). We used a linear regression model accounting for biological and technical covariates depending upon the cohort to be analyzed. The final model used was implemented in R version 3.6.1 (R Core Team, 2019) is as follows:

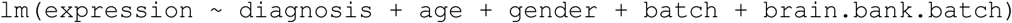

#### Consensus WGCNA

Consensus weighted co-expression network analysis (cWGCNA) was performed in R (R Core Team, 2019) using WGCNA package (Langfelder and Horvath, 2008). To identify consensus proteomics modules across different brain banks, we employed a signed cWGCNA approach by calculating component-wise values for topological overlap for individual brain banks. Biweighted mid-correlations were calculated for all pairs of genes, and then a signed similarity matrix was created. In the signed network, the similarity between genes reflects the sign of the correlation of their expression profiles. The signed similarity matrix was then raised to power β to emphasize strong correlations and reduce the emphasis of weak correlations on an exponential scale. The resulting adjacency matrix was then transformed into a topological overlap matrix (TOM). The consensus TOM (cTOM) was calculated by taking the component-wise minimum of TOMs in each dataset. Using 1 − cTOM as the distance measure, genes were hierarchically clustered (Parikshak et al., 2016). Module assignments were determined using a dynamic tree-cutting algorithm (cutreeHybrid, using the default parameters except deepSplit = 4, cutHeight = 0.999, minModuleSize = 20, dthresh = 0.07 and pamStage = FALSE). Network visualization was performed with the igraph package in R (Csardi and Nepusz, 2006).

Module eigenproteins were calculated as the first principal component of the coexpressed genes in the module (Langfelder and Horvath, 2007; Zhang and Horvath, 2005). The eigengene-based connectivity (kME) was used to represent the strength of a gene’s correlation with other gene module members. It is defined as the correlation between a gene’s expression and the module eigengene (Langfelder and Horvath, 2007; Zhang and Horvath, 2005). Hub genes of a module were considered genes with high kME.

### Quantification and Statistical Analysis

#### Cell-type and Gene-ontology enrichment

For gene-set enrichment analysis of cell type (Zhang et al., 2016) and gene ontology (Ashburner et al., 2000; Gene Ontology Consortium, 2019) we use the hypergeometric test, which is equivalent to a one-sided Fisher’s exact test, to evaluate whether a gene set is enriched over background, providing a p-value and enrichment value for gene set enrichment (Parikshak et al., 2016). An assumption with such a test is that the background set is similar to the gene set considered for enrichment in all factors other than pathway membership.

#### Module Eigenprotein Association with Disease

We determined if the module eigenprotein was significantly different from disease diagnosis (AD or AsymAD) and control using the Wilcoxon rank sum test (Wilcoxon, 1945). Module eigenproteins were correlated with clinical score (CASI) and neuropathological scores (CERAD, Braak) using Pearson correlation (Pearson, 1931). We used the same method for clinical scores and neuropathological scores in UPenn and Emory datasets. For UPenn and Emory, we determined if the module eigenprotein was significantly different from disease diagnosis and control using linear regression.

#### Comparison of the AD proteome and transcriptome

We performed weighted co-expression network analysis (WGCNA) (Langfelder and Horvath, 2008) for the Mayo proteomic and transcriptomic data resulting in a proteomic network and transcriptomic network. Biweighted mid-correlations were calculated for all pairs of genes, and then a signed similarity matrix was created. The signed similarity matrix was then raised to power β to emphasize strong correlations and reduce the emphasis of weak correlations on an exponential scale. The resulting adjacency matrix was then transformed into a topological overlap matrix (TOM). Using 1 − TOM as the distance measure, genes were hierarchically clustered (Parikshak et al., 2016).

We identified consensus proteomic modules overlapping with Mayo proteomic modules, and Mayo proteomic modules overlapping with Mayo transcriptomic modules using the Fisher’s exact test (Fisher, 1922).

#### Differential expression analysis in PSP compared to AD

We used the set of differentially expressed genes in PSP compared to AD (FDR<0.1) at the RNA and protein level. We correlated the fold-change in RNA level to the fold-change in protein level with linear regression and Wald test to determine significance. We performed gene ontology enrichment of genes upregulated and downregulated in AD compared to PSP at both the RNA and protein levels using the Database for Annotation, Visualization and Integrated Discovery (DAVID) (Huang et al., 2009).

#### Module Connectivity in Proteomic versus Transcriptomic Modules

We compared intersecting genes from Mayo proteomic modules that had significant hypergeometric overlap with Mayo transcriptomic modules. To calculate the shortest path between all pairs of genes in a module, we first removed all edges with correlations from the adjacency matrix lower than the 75^th^ percentile. The shortest path was calculated with the igraph package in R (Csardi and Nepusz, 2006). For pairs of genes that were connected, we determined if the mean shortest path in the Mayo proteomic module was significantly different than the mean shortest path in the Mayo transcriptomic modules using Student’s t-test (Student, 1908). We calculated the clustering coefficient, which measures the connectivity among a gene’s neighbors, using the WGCNA R package (Langfelder and Horvath, 2008) and determined if it was significantly different in Mayo proteomic modules versus transcriptomic modules using Student’s t-test (Student, 1908).

#### GWAS Enrichment

GWAS summary data was obtained from the International Genomics of Alzheimer’s Project (IGAP) for AD (Lambert et al., 2013) and PSP (Höglinger et al., 2011). For this analysis, we used the MAGMA approach (de Leeuw et al., 2015) to assign each gene a score based on the best p-value of a SNP in a given GWAS study within 20kB of the gene, and then set a p-value cut-off at 0.05 to define the gene as included in the common variant set related to that study. We performed enrichment analysis with logistic regression controlling for gene length and other biases (Parikshak et al., 2016).

### Data and Code Availability

All the proteomics data is available via synapse link: https://www.synapse.org/#!Synapse:syn2580853/wiki/409840 Code is available on GitHub: https://github.com/dhglab/Identification-of-conserved-proteomic-networks-in-neurodegenerative-dementia

### Additional Resources

The following website (https://coppolalab.ucla.edu/gclabapps/nb/browser?id=SwarupConsensusProteomicsV4) includes an interactive graphical interface for the consensus proteomic network, allowing exploration of individual proteins and their relationships.

## Supplemental Information

Figure S1. Related to Figure 2. Consensus Proteomics Analyses

Consensus proteomics dendrogram showing the proteomics modules from 5 different proteomics datasets. Color bars below the modules give information on correlation of disease condition (AD=Alzheimer’s disease, Poss AD=possible AD, Prob AD=probable AD, Def AD=definite AD, Asym AD=asymptomatic AD, PSP= Progressive Supranuclear Palsy, MCI=mild cognitive impairment) and biological covariates (age and gender) with the expression of a particular gene. Red is positive correlation, and blue is anti-correlation. cTOM=consensus Topological Overlap Matrix, MSBB=Mount Sinai Brain Bank, ACT=Adult Changes of Thought, BLSA= Baltimore Longitudinal Study of Aging

Figure S2. Related to Figure 3. Early proteomic changes in AD.

(A-I) Plots showing C8 module eigenprotein trajectory with diagnosis (A) and CDR (B) in the MSBB dataset; diagnosis (C) in the Mayo dataset; diagnosis (D), CERAD (E) and BRAAK (F) in the BLSA; and diagnosis (G), CERAD (H) and BRAAK (I) in the Banner dataset. (J-R) Plots showing C8 module eigenprotein trajectory with diagnosis (J) and CDR (K) in the MSBB dataset; diagnosis (L) in the Mayo dataset; diagnosis (M), CERAD (N) and BRAAK (O) in the BLSA; and diagnosis (P), CERAD (Q) and BRAAK (R) in the Banner dataset. MSBB=Mount Sinai Brain Bank, ACT=Adult Changes of Thought, BLSA= Baltimore Longitudinal Study of Aging, AD=Alzheimer’s disease, Poss AD=possible AD, Prob AD=probable AD, Def AD=definite AD, Asym AD=asymptomatic AD, MCI=mild cognitive impairment, crit not met=criteria not met. *p<0.05; **p<0.01; ***p<0.005; ns=non significant

Figure S3. Related to Figure 4. Late proteomic changes for C1 and C2 in AD.

(A-L) Plots showing C1 module eigenprotein trajectory with diagnosis (A) and CDR (B) in the MSBB dataset; diagnosis (C) in the Mayo dataset; diagnosis (D), CERAD (E) and BRAAK (F) in the BLSA; and diagnosis (G), CERAD (H) and BRAAK (I) in the Banner dataset. (J-L) Validation of C1 module trajectory using Emory dataset (Ping et al 2018) showing the eigenprotein trajectory with diagnosis (J), CERAD (K) and BRAAK score (L). (M-X) Plots showing C2 module eigenprotein trajectory with diagnosis (M) and CDR (N) in the MSBB dataset; diagnosis (O) in the Mayo dataset; diagnosis (P), CERAD (Q) and BRAAK (R) in the BLSA; and diagnosis (S), CERAD (T) and BRAAK (U) in the Banner dataset. (V-X) Validation of C2 module trajectory using Emory dataset (Ping et al 2018) showing the eigenprotein trajectory with diagnosis (V), CERAD (W) and BRAAK score (X). AD=Alzheimer’s disease, Poss AD=possible AD, Prob AD=probable AD, Def AD=definite AD, Asym AD=asymptomatic AD, MCI=mild cognitive impairment, crit not met=criteria not met, ADPD=Alzheimer’s disease with Parkinson’s disease, PD=Parkinson’s disease. *p<0.05; **p<0.01; *** p<0.005; n.s.=non-significant

Figure S4. Related to Figure 4. Late proteomic changes for C3 and C7 in AD.

(A-L) Plots showing C3 module eigenprotein trajectory with diagnosis (A) and CDR (B) in the MSBB dataset; diagnosis (C) in the Mayo dataset; diagnosis (D), CERAD (E) and BRAAK (F) in the BLSA; and diagnosis (G), CERAD (H) and BRAAK (I) in the Banner dataset. (J-L) Validation of C3 module trajectory using Emory dataset (Ping et al 2018) showing the eigenprotein trajectory with diagnosis (J), CERAD (K) and BRAAK scores (L). (M-X) Plots showing C7 module eigenprotein trajectory with diagnosis (M) and CDR (N) in the MSBB dataset; diagnosis (O) in the Mayo dataset; diagnosis (P), CERAD (Q) and BRAAK (R) in the BLSA; and diagnosis (S), CERAD (T) and BRAAK (U) in the Banner dataset. (V-X) Validation of C7 module trajectory using Emory dataset (Ping et al 2018) showing the eigenprotein trajectory with diagnosis (V), CERAD (W) and BRAAK scores (X). AD=Alzheimer’s disease, Poss AD=possible AD, Prob AD=probable AD, Def AD=definite AD, Asym AD=asymptomatic AD, MCI=mild cognitive impairment, crit not met=criteria not met, ADPD=Alzheimer’s disease with Parkinson’s disease, PD=Parkinson’s disease. *p<0.05; **p<0.01; ***p<0.005; n.s.=non-significant

Figure S5. Related to Figure 5. Mayo Proteomic Module Analysis

Multidimensional scaling plot demonstrates relationship between Mayo proteomics modules and clustering by cell-type relationship. Also shown are the major Gene Ontology enrichment, cell-type enrichment, and hubs for that module. Modules upregulated in AD are on the left and modules downregulated in AD are on the right.

Figure S6. Related to Figure 5. Hypergeometric Overlap

(A) Hypergeometric overlap of consensus proteomics and Mayo proteomics modules. (B) Hypergeometric overlap of Mayo proteomics and Mayo transcriptomic modules. Values represents odds ratio. Color bar represents −log10 p-value of enrichment. *p<0.05; **p<0.01; ***p<0.005

Table S1. Related to STAR Methods section “Human Samples”. Meta-data from BLSA, ACT, MSSBB, Banner, Mayo and UPenn datasets

Table S2. Related to Results section “Identification of robust disease-relevant protein co-expression signature”. Consensus Proteomic Module Membership

