## Supplemental Information for "Identification of conserved proteomic networks in neurodegenerative dementia"

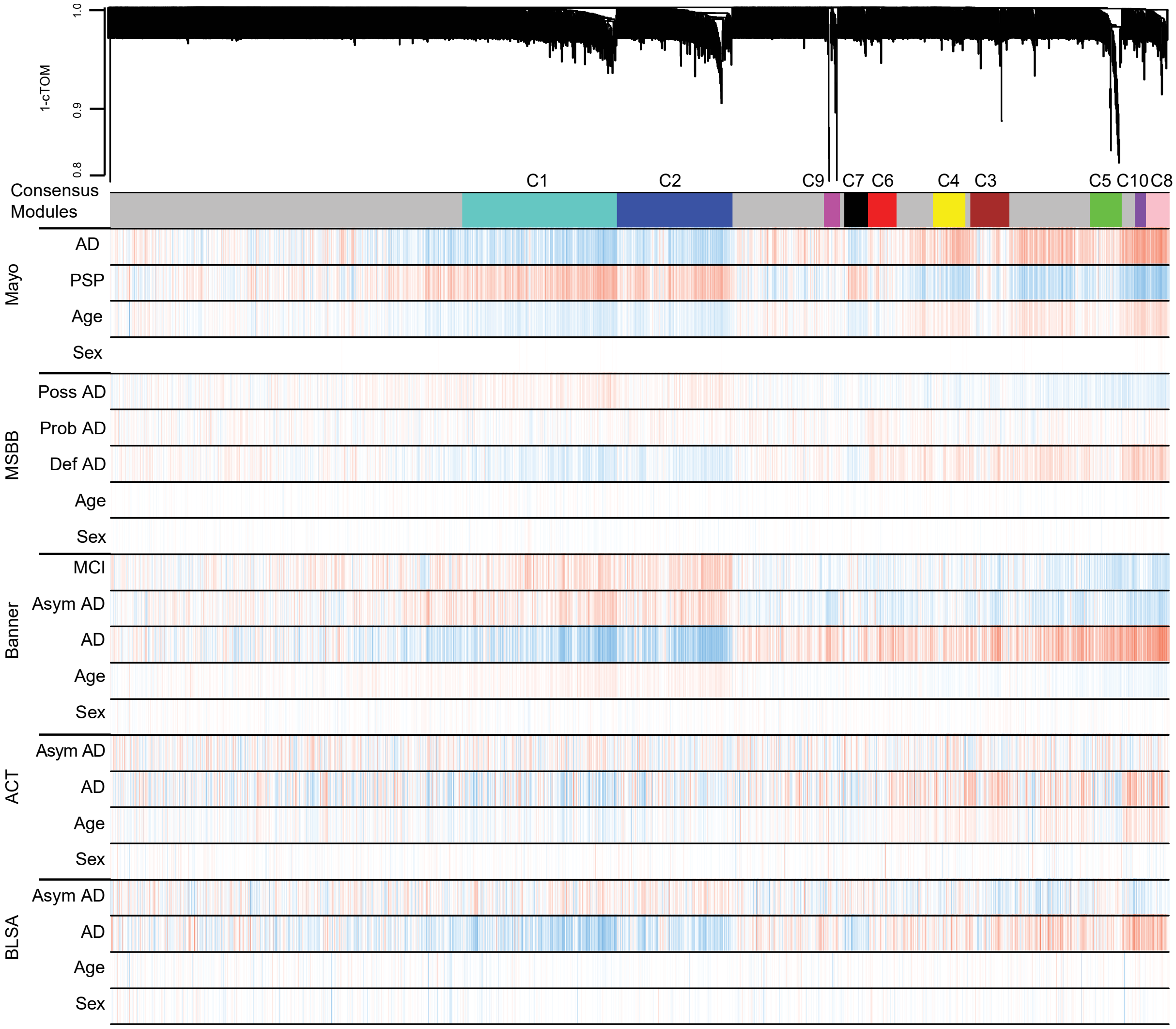


Figure S1. Related to Figure 2. Consensus Proteomics Analyses
Consensus proteomics dendrogram showing the proteomics modules from 5 different proteomics datasets. Color bars below the modules give information on correlation of disease condition (AD=Alzheimer's disease, Poss AD=possible AD, Prob AD=probable AD, Def AD=definite AD, Asym AD=asymptomatic AD, PSP= Progressive Supranuclear Palsy, MCI=mild cognitive impairment) and biological covariates (age and gender) with the expression of a particular gene. Red is positive correlation, and blue is anti-correlation. cTOM=consensus Topological Overlap Matrix, MSBB=Mount Sinai Brain Bank, ACT=Adult Changes of Thought, BLSA= Baltimore Longitudinal Study of Aging

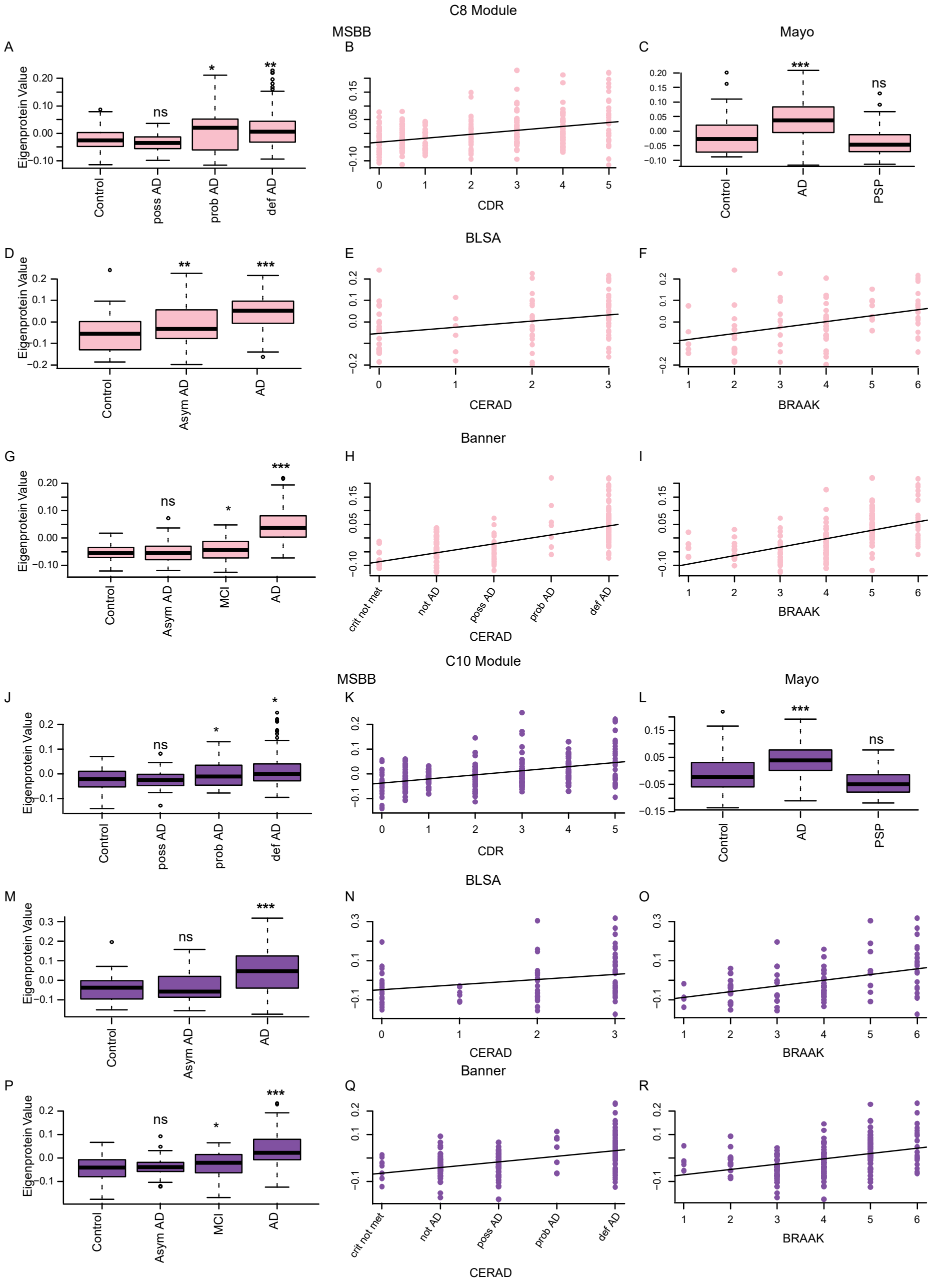


Figure S2. Related to Figure 3. Early proteomic changes in AD.
(A-I) Plots showing C8 module eigenprotein trajectory with diagnosis (A) and CDR (B) in the MSBB dataset; diagnosis (C) in the Mayo dataset; diagnosis (D), CERAD (E) and BRAAK (F) in the BLSA; and diagnosis (G), CERAD (H) and BRAAK (I) in the Banner dataset. (J-R) Plots showing C8 module eigenprotein trajectory with diagnosis (J) and CDR (K) in the MSBB dataset; diagnosis (L) in the Mayo dataset; diagnosis (M), CERAD (N) and BRAAK (O) in the BLSA; and diagnosis (P), CERAD (Q) and BRAAK (R) in the Banner dataset. MSBB=Mount Sinai Brain Bank, ACT=Adult Changes of Thought, BLSA= Baltimore Longitudinal Study of Aging, AD=Alzheimer’s disease, Poss AD=possible AD, Prob AD=probable AD, Def AD=definite AD, Asym AD=asymptomatic AD, MCI=mild cognitive impairment, crit not met=criteria not met. *p<0.05; **p<0.01; ***p<0.005; ns=non significant

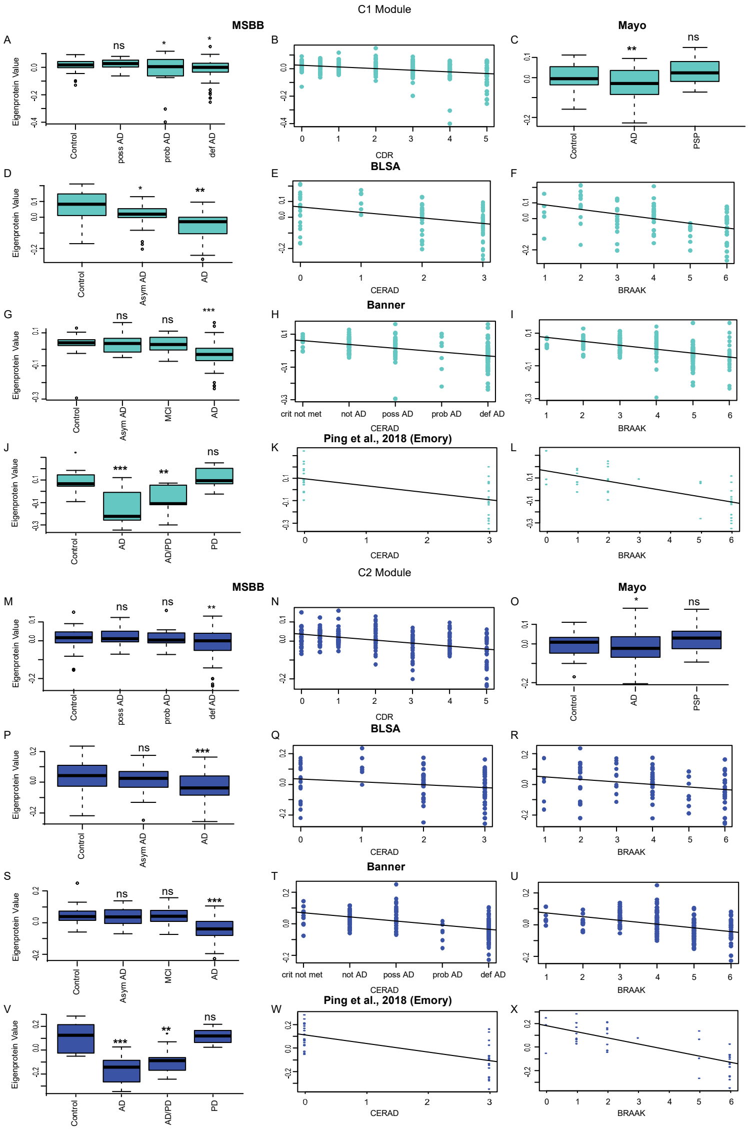


Figure S3. Related to Figure 4. Late proteomic changes for C1 and C2 in AD.
(A-L) Plots showing C1 module eigenprotein trajectory with diagnosis (A) and CDR (B) in the MSBB dataset; diagnosis (C) in the Mayo dataset; diagnosis (D), CERAD (E) and BRAAK (F) in the BLSA; and diagnosis (G), CERAD (H) and BRAAK (I) in the Banner dataset. (J-L) Validation of C1 module trajectory using Emory dataset (Ping et al 2018) showing the eigenprotein trajectory with diagnosis (J), CERAD (K) and BRAAK score (L). (M-X) Plots showing C2 module eigenprotein trajectory with diagnosis (M) and CDR (N) in the MSBB dataset; diagnosis (O) in the Mayo dataset; diagnosis (P), CERAD (Q) and BRAAK (R) in the BLSA; and diagnosis (S), CERAD (T) and BRAAK (U) in the Banner dataset. (V-X) Validation of C2 module trajectory using Emory dataset (Ping et al 2018) showing the eigenprotein trajectory with diagnosis (V), CERAD (W) and BRAAK score (X). MSBB=Mount Sinai Brain Bank, ACT=Adult Changes of Thought, BLSA= Baltimore Longitudinal Study of Aging, AD=Alzheimer's disease, Poss AD=possible AD, Prob AD=probable AD, Def AD=definite AD, Asym AD=asymptomatic AD, MCI=mild cognitive impairment, crit not met=criteria not met, ADPD=Alzheimer's disease with Parkinson's disease, PD=Parkinson's disease. *$p<0.05$; **$p<0.01$; *** $p<0.005$; n.s.=non-significant

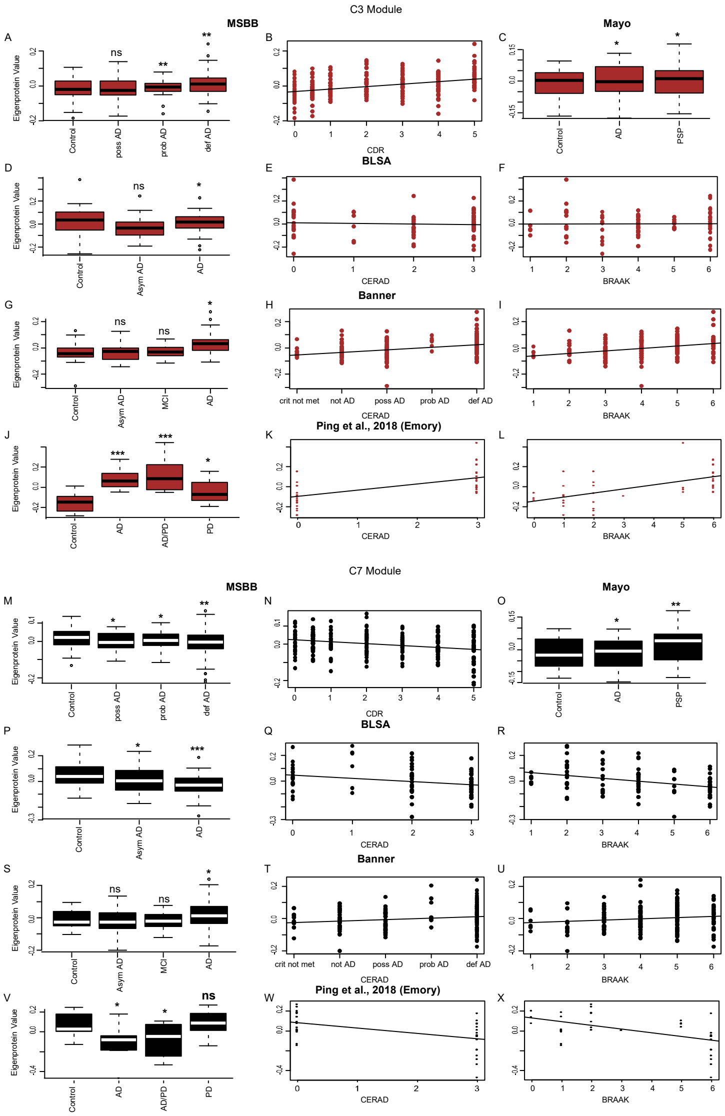


Figure S4. Related to Figure 4. Late proteomic changes for C3 and C7 in AD.
(A-L) Plots showing C3 module eigenprotein trajectory with diagnosis (A) and CDR (B) in the MSBB dataset; diagnosis (C) in the Mayo dataset; diagnosis (D), CERAD (E) and BRAAK (F) in the BLSA; and diagnosis (G), CERAD (H) and BRAAK (I) in the Banner dataset. (J-L) Validation of C3 module trajectory using Emory dataset (Ping et al 2018) showing the eigenprotein trajectory with diagnosis (J), CERAD (K) and BRAAK scores (L). (M-X) Plots showing C7 module eigenprotein trajectory with diagnosis (M) and CDR (N) in the MSBB dataset; diagnosis (O) in the Mayo dataset; diagnosis (P), CERAD (Q) and BRAAK (R) in the BLSA; and diagnosis (S), CERAD (T) and BRAAK (U) in the Banner dataset. (V-X) Validation of C7 module trajectory using Emory dataset (Ping et al 2018) showing the eigenprotein trajectory with diagnosis (V), CERAD (W) and BRAAK scores (X). MSBB=Mount Sinai Brain Bank, ACT=Adult Changes of Thought, BLSA= Baltimore Longitudinal Study of Aging, AD=Alzheimer's disease, Poss AD=possible AD, Prob AD=probable AD, Def AD=definite AD, Asym AD=asymptomatic AD, MCI=mild cognitive impairment, crit not met=criteria not met, ADPD=Alzheimer's disease with Parkinson's disease, PD=Parkinson's disease. *p<0.05; **p<0.01; ***p<0.005; n.s.=non-significant

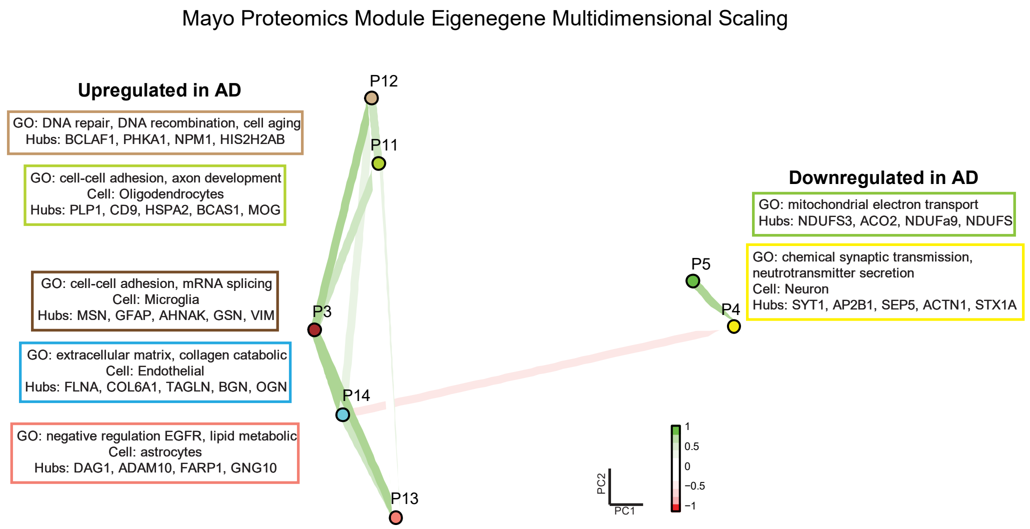


Figure S5. Related to Figure 5. Mayo Proteomic Module Analysis

Multidimensional scaling plot demonstrates relationship between Mayo proteomics modules and clustering by cell-type relationship. Also shown are the major Gene Ontology enrichment, cell-type enrichment, and hubs for that module. Modules upregulated in AD are on the left and modules downregulated in AD are on the right.

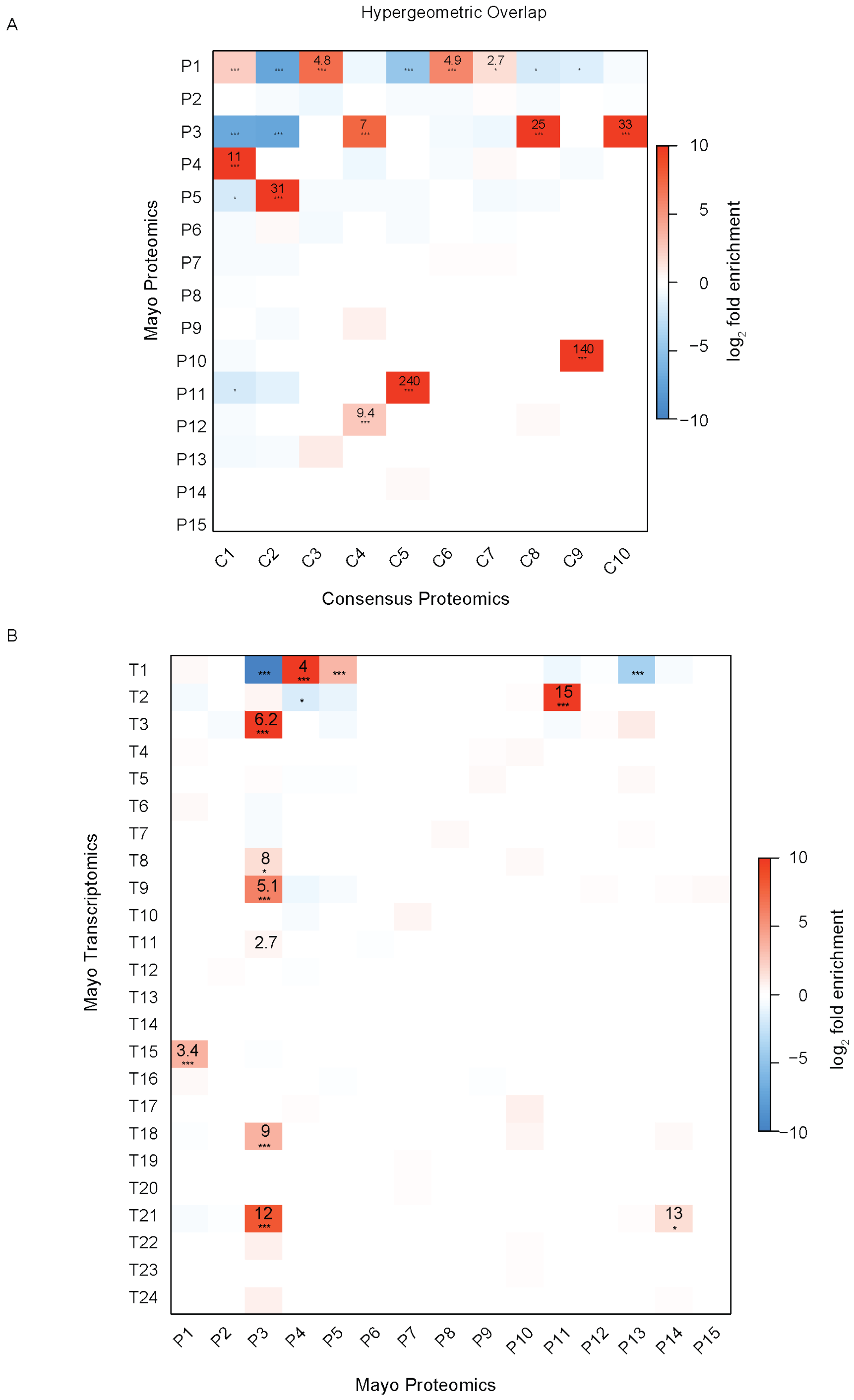


Figure S6. Related to Figure 5. Hypergeometric Overlap
(A) Hypergeometric overlap of consensus proteomics and Mayo proteomics modules. (B) Hypergeometric overlap of Mayo proteomics and Mayo transcriptomic modules. Values represents odds ratio. Color bar represents -log10 p-value of enrichment. *p<0.05; **p<0.01; ***p<0.005

| Module | AD stage | AD Correlation | Gene Ontology | Cell Type | Hubs | Other Disease Trajectory | |
|---|---|---|---|---|---|---|---|
| | | | | | | UPenn | Emory |
| C8 | early | ↑ | cell-adhesion, immune response | astrocytes | GFAP, MSN, GNAI3, DDAH2 | ↑ AD, FTD TDP, PSP-CBD, PD-D | ↑ AD, ADPD |
| C10 | early | ↑ | cell-cell contact, cell-adhesion | microglia | VIM, ANXA1, PLCD1, AHNAK | ↑ AD, FTD TDP, PSP-CBD | ↑ AD, ADPD |
| C1 | late | ↓ | synaptic process, neurotransmitter secretion | neuron | SYT1, HOMER1, STX1A, ATP6V0D1 | ↓ AD, FTD TDP | ↓ AD, ADPD |
| C2 | late | ↓ | mitochondria electron transport chain | GABAergic Neuron | NDUFS1, NDUFA10, ACO2, | ↓ AD, FTD TDP, PSP-CBD, PD-D, MSA, ALS, PD | ↓ AD, ADPD |
| C3 | late | ↑ | MAPK signaling, Fc-receptor signaling, NF-kB signaling | | VCP, MAPK1, MAPK3, NCAM2 | ↑ AD, FTD TDP, PSP-CBD, PD-D | ↑ AD, ADPD, PD |
| C7 | late | ↓ | protein localization and transport | | TUBB4B, CCTs, DCTN1 | ↑ AD, FTD TDP, PSP-CBD | ↓ AD, ADPD |

Table S3. Related to Results section “Identification of robust disease-relevant protein co-expression signature.” Consensus Proteomics Module Summary. AD=Alzheimer’s disease, FTD TDP=Frontotemporal dement with TDP-43, PSP-CBD=Progressive Supranuclear Palsy with Corticobasal Degeneration, PD=Parkinson’s disease, PD-D=Parkinson’s disease with, MSA=Multiple Systems Atrophy, ALS=Amyotrophic Lateral Sclerosis, ADPD=Alzheimer’s disease with Parkinson’s disease

| Protein, Trans | Module Size (protein, trans) | # Intersecting Genes | Intermodular Connectivity (kIM) | | | Shortest Path # not connected | | Clustering Coefficient for intersecting genes | | |
|---|---|---|---|---|---|---|---|---|---|---|
| | | | Protein (mean,sd) | Trans (mean, sd) | t test p-value | Protein | Trans | Protein (mean,sd) | Trans (mean, sd) | t test p-value |
| P3, T3 | (335 , 431) | 55 | 0.72 ( 0.8 ) | 10.5 (2.8) | <2.2E-16 | 107 | 0 | 0.018 (0.01) | 0.064 (0.006) | <2.2E-16 |
| P3, T8 | (335 , 302) | 24 | 0.37 ( 0.32 ) | 2.13 ( 0.78 ) | 9.05E-11 | 0 | 0 | 0.021 (0.01) | 0.044 (0.005) | 9.10E-11 |
| P3, T9 | (335 , 307) | 6 | 0.02 ( 0.01 ) | 1.25 ( 0.42 ) | 7.50E-04 | 0 | 0 | 0.011 (0.003) | 0.056 (0.016) | 7.20E-04 |
| P3, T18 | (335 , 156) | 12 | 0.25 ( 0.24 ) | 1.2 ( 0.41 ) | 5.40E-06 | 11 | 0 | 0.026 (0.01) | 0.044 (0.004) | 8.80E-04 |
| P3, T21 | (335 , 137) | 18 | 0.33 ( 0.27 ) | 4.23 ( 0.77 ) | 2.50E-13 | 33 | 0 | 0.024 (0.01) | 0.058 (0.004) | 8.80E-11 |
| P4, T1 | (281, 3698) | 177 | 2.74 ( 2.72 ) | 26.28 ( 14.61 ) | <2.2E-16 | 2046 | 176 | 0.023 ( 0.009 ) | 0.11 (0.028) | <2.2E-16 |

Table S4. Related to Results section "Comparison of the AD proteome and transcriptome". Mayo proteomic and transcriptomic module connectivity.
